# Multiplexed Detection of Molecular Interactions with DNA Origami Engineered Cells in 3-D Collagen Matrices

**DOI:** 10.1101/2022.02.04.479158

**Authors:** Melika Shahhosseini, Peter E. Beshay, Ehsan Akbari, Alex Avendano, Jonathan W. Song, Carlos E. Castro

**Affiliations:** Department of Mechanical and Aerospace Engineering, The Ohio State University, Columbus, OH, 43210, United States of America; Biophysics Graduate Program, The Ohio State University, Columbus, OH, 43210, United States of America; Department of Biomedical Engineering, The Ohio State University, Columbus, OH, 43210, United States of America; Comprehensive Cancer Center, The Ohio State University, Columbus, OH, 43210 United States of America

**Keywords:** DNA origami, DNA nanotechnology, subcellular interactions, cell membrane engineering, cell microenvironment

## Abstract

The interactions of cells with signaling molecules present in their local microenvironment maintain cell proliferation, differentiation, and spatial organization, and mediate progression of diseases such as metabolic disorders and cancer. Real-time monitoring of the interactions between cells and their extracellular ligands in a 3-D microenvironment can inform detection and understanding of cell processes and development of effective therapeutic agents. DNA-origami technology allows for the design and fabrication of biocompatible and 3-D functional nanodevices via molecular self-assembly for various applications including molecular sensing. Here, we report a robust method to monitor live cell interactions with molecules in their surrounding environment in a 3-D tissue model using a microfluidic device. We used a DNA origami Cell Sensing Platform (CSP) to detect two specific nucleic acid sequences on the membrane of B cells and dendritic cells. We further demonstrated real-time detection of biomolecules with the DNA sensing platform on the surface of dendritic cells in a 3-D microfluidic tissue model. Our results establish the integration of live cells with membranes engineered with DNA nanodevices into microfluidic chips as a highly capable biosensor approach to investigate subcellular interactions in physiologically relevant 3-D environments under controlled biomolecular transport.

## 1. Introduction

Living cells interact with a myriad of extracellular signaling molecules, such as nucleic acids, growth factors, and cytokines. For instance, antigen-presenting dendritic cells (DC) are activated by cytokines in the local environment, leading to their secretion of pro-inflammatory cytokines, activation of T-cells, and regulation of immune response and homeostasis^[1]^. DCs also respond to extracellular microRNAs (miRNAs) and tumor-secreted DNA, to modulate immune responses in cancer and other diseases^[2–4]^. Yet, our understanding of interactions between cells and extracellular ligands, especially in native 3-D tissue environments, remains incomplete. Many cellular responses to external cues occur rapidly and are contingent on the spatial context of the signals (e.g., molecular gradients) within the local milieu. Therefore, improvements in cell-based biosensor technologies with dynamic subcellular readouts and improved spatiotemporal resolution are needed to further our understanding of cell interactions with extracellular signaling molecules in their microenvironment. Thus, the goal of this work is to develop a robust approach to monitor cell interactions with the surrounding environment in real-time in a physiologically relevant 3-D microenvironment, which would enhance studies focused on understanding cell biology and developing effective therapeutics.

Many of the conventional molecular biochemistry assays to characterize cell interactions, such as flow cytometry, enzyme-linked immunosorbent assay (ELISA), immunostaining, and polymerase chain reaction (PCR), focus on measuring the presence of specific molecular markers that are indicative of a cell response (e.g. membrane receptors, mRNA, or activated signaling molecules). However, they require staining, washing, and manipulation before imaging and only provide endpoint results. These methods are further constrained by requiring the measurement of many cells as opposed to single cell analysis. They also, do not provide a direct indication of the extracellular ligand that leads to cell activation (only measure cell response), and hence, fail to monitor interactions at the cell-surface in real-time. More recent studies have engineered cell surfaces with synthetic probes^[5,6]^, labeled proteins^[7]^, or fluorescent DNA constructs^[8]^. These advances have allowed for improved capabilities for measurements at the single cell level^[9]^, but they still are only capable of understanding cell interactions with a single target molecule and cannot monitor the interaction of cell membranes with multiple biomolecules.

Cell interactions are highly affected by their microenvironments, including the physical and biochemical properties of the surrounding extracellular matrix (ECM)^[10,11]^. Living systems are complex and difficult to control and interrogate efficiently at cellular length scales making it difficult to study detailed mechanisms efficiently in animal models. 3-D tissue models are useful surrogates for animal models that recapitulate key aspects of living tissues while enabling control over the environment with simpler measurement readouts; hence these are useful systems to study underlying mechanisms of cell response in physiologically relevant microenvironment. Although prior work has demonstrated that engineered cells with sensing capabilities can be monitored in live animal models^[5,8]^, they have not yet been widely deployed in reconstituted 3-D tissue models, which provide simpler measurement readouts and control over the microenvironment composition. The incorporation of cells engineered with molecular detection capabilities into 3-D tissue models would enable novel insights into the spatiotemporal effects of soluble signals and intercellular communication, including mechanisms that mediate disease progression.

Structural DNA nanotechnology^[12]^ has emerged as a versatile approach to make biocompatible nanodevices with precise structure that can be functionalized with a large range of molecules, making them attractive for biological applications, including engineering cell membranes^[13]^. The molecular self-assembly process known as DNA-origami^[14]^ allows for programming complex nanoscale geometry ^[15,16]^, tunable mechanical and dynamic properties^[17–19]^, and the incorporation of one or many molecules with nanometer precision ^[20–25]^. DNA origami nanodevices have been recently used in applications including drug delivery^[26–29]^, ion and molecular transport^[30,31]^, imaging^[32]^, as well as molecular sensing, manipulation, and measurement^[21,33,34]^. In addition, 3-D DNA nanodevices were successfully incorporated into the cell membranes to control adhesion between two living cells^[20]^, and facilitate cell-cell communication^[35]^, and other advanced functions like membrane sculpting^[36]^ or cargo transport^[37]^ have been demonstrated on synthetic membranes. While these nanodevices have been used in a variety of biological assays including cell culture ^[29,38]^, cell spheroids^[39]^, and animal models^[27]^, they have not been implemented into 3-D ECM model systems, which are ideal for probing biological mechanisms in native tissue environments.

Here, we establish a method to sense multiple biomolecules on the membrane of living cells in a 3D tissue model. We designed a DNA origami Cell Sensing Platform (CSP) capable of detecting the presence of two specific molecules. We focused on detecting nucleic acid sequences on the surfaces of both CH12-LX B cells (suspension) and MutuDC 1949 dendritic cells (adherent) with fluorescence-based reporting both in cell culture and in 3-D collagen matrices. Using microfluidics to control the ECM structure formation and the localized transport of target molecules, we show how multifunctional DNA origami devices can be used to probe the temporal interactions of cells and their local environment with subcellular resolution in a tissue model system.

## 2. Results and Discussion

### 2.1: A stable DNA origami device detects multiple targets

We designed the CSP structure using the DNA design software caDNAno^[40]^. The structure consists of 40 double-stranded DNA (dsDNA) helices organized into three layers with gaps in the middle layer (Figure 1, A), inspired by a prior nanorod design that exhibited efficient folding and robust stability in cell culture media^[29]^. The full caDNAno design and corresponding oligonucleotide sequences are provided in Figure S1 and Table S1. The CSP design allows for the selective incorporation of up to 30 single-stranded DNA (ssDNA) overhangs on the membrane-facing side (bottom overhangs) and up to 12 overhangs on the membrane-opposing side (top overhangs) (Figure 1, A and D). To ensure proper molecular self-assembly and optimize the ion concentration and isothermal annealing temperature ^[41]^, CSP was subjected to thermal annealing by rapid heating to 65 °C followed by slow cooling to 4 °C over 2.5 d in different MgCl_2_ concentrations (10 to 24 mM). Gel electrophoresis confirmed folding of CSP in a wide range of 14 to 24 mM MgCl_2_ (Figure S2, A). We chose 18mM for subsequent folding. Self-assembled nanostructures were then folded at various isothermal annealing temperatures in 18mM MgCl_2_ over 4 hours and subjected to agarose gel electrophoresis. Gel electrophoresis revealed the successful folding of CSP over the annealing temperature range of 42 to 59°C (Figure S2, B). We chose 52°C for subsequent folding with isothermal annealing. To ensure that the structural integrity is preserved in cell culture media, we tested the stability of CSP in RPMI 1640 cell culture medium or 1 x Phosphate Buffered Saline (PBS) supplemented with 2% or 8% Fetal Bovine Serum (FBS) and 1mM MgCl_2_ after 4 hours incubation in 37 °C via agarose gel electrophoresis. Figure 1, B shows that CSP nanostructures remained intact under each cell culture media conditions compared to the control structures in storage buffer, 1x FOB solution (5 mM Tris, 5 mM NaCl, 1 mM EDTA) supplemented with 10 mM MgCl_2_. The slight differences in the migration speed is likely due to the different buffer conditions. Leading bands were excised and visualized using transmission electron microscopy (TEM) to confirm that CSP preserved its structural integrity in cell culture media (Figure 1, C and Figure S3).

**Figure 1:**
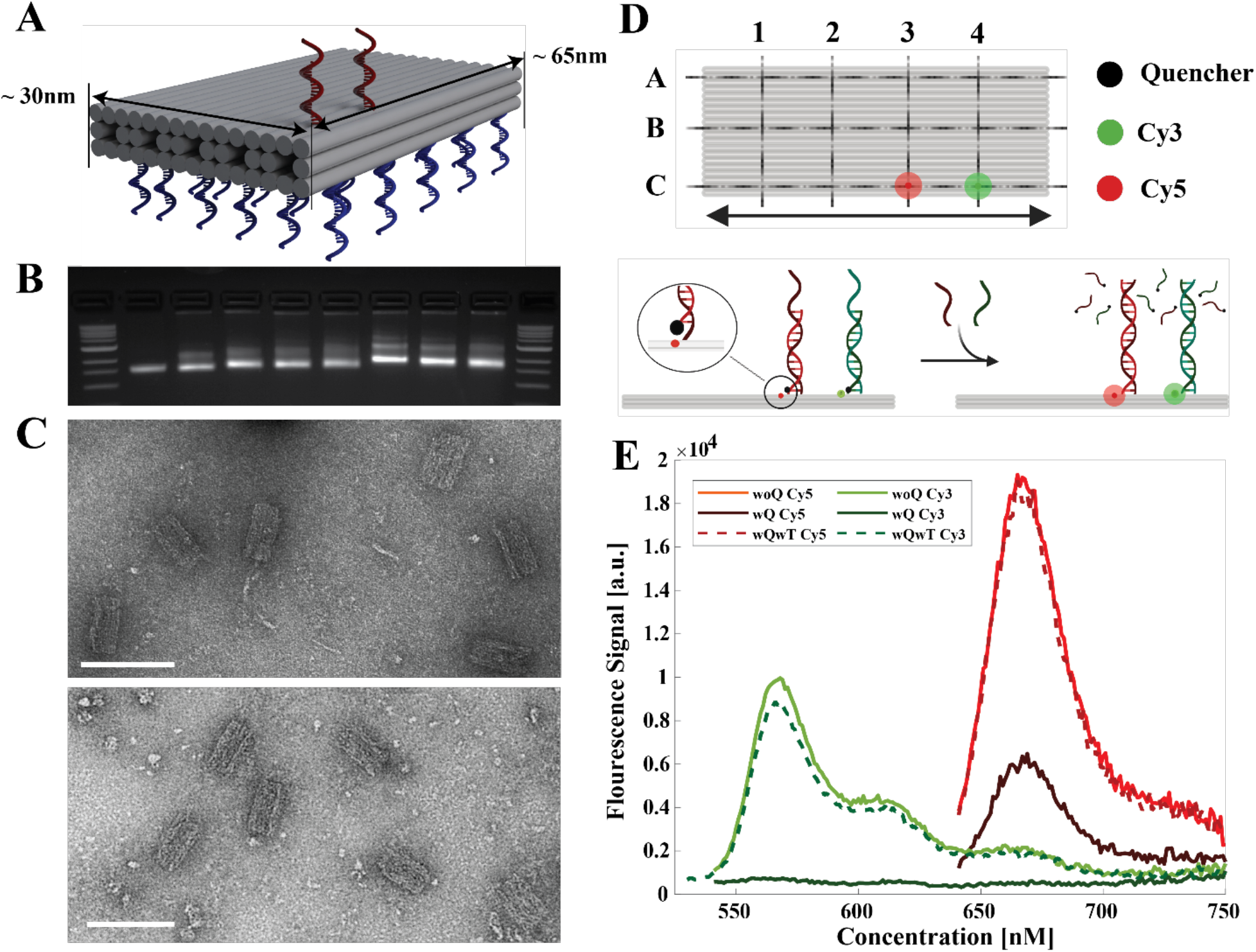
Design and Characterization of CSP. A) Isometric schematic illustrating the structure design and locations of the top (red) and bottom (blue) overhangs. B) Gel analysis confirms CSP structural stability after 4 h in cell culture conditions (37 °C) compared to CSP in storage buffer (left to right, 1 kb DNA ladder, the 7249 M13mp18 scaffold starting material, CSP in storage buffer, CSP in RPMI supplemented with 1 mM MgCl_2_, CSP in RPMI supplemented with 1 mM MgCl_2_ and 2% FBS, CSP in RPMI supplemented with 1 mM MgCl_2_ and 8% FBS, CSP in 1xPBS, CSP in 1xPBS supplemented with 1 mM MgCl_2_ and 2% FBS, CSP in 1xPBS supplemented with 1 mM MgCl_2_ and 8% FBS, 1 kb DNA ladder). C) TEM images confirm the structure is preserved after incubation under cell culture conditions for 4 h in storage buffer (top) and RPMI supplemented with 2% FBS and 1mM MgCl_2_ (bottom). Scale bars: 100nm. D) 2D top-view and side-view schematics of CSP with DNA detection modules on C3 (red) and C4 (green) locations. Before the introduction of DNA targets, the quenching oligo is bound to the respective top overhangs, placing the quencher in the proximity of fluorophore, leading to a low fluorescent signal. After the addition of DNA targets, the quenching oligo is displaced with the target oligo, resulting in a high fluorescent signal (shown in E, dashed lines). E) Ensemble fluorescent measurement showing the fluorescent signal of CSP folded with quenching oligo (dark solid lines), folded without the quenching oligo (light solid lines) and folded with quenching oligo followed by the addition of target oligo (dashed lines).

The CSP enables incorporation of up to 12 top overhangs (Figure 1, D) which could potentially allow for 12 distinct detection sites for various biomolecular targets. Here, to enable the detection of two distinct DNA target strands, two overhangs with unique sequences were incorporated at locations 3C and 4C onto the CSP. Two quencher-labeled oligonucleotides (QO) complementary to each of the top overhangs were designed to provide a 5-base pair toehold for the binding of target oligos (Figure 1, D). Moreover, one Cy3-labeled oligonucleotide and one Cy5-labeled oligonucleotide were incorporated into the CSP platform to create the two-channel internally labeled CSP. The Cy3 and Cy5 oligonucleotides were incorporated so that the Cy3 molecule is proximal to the quencher at the 4C overhang location, and the Cy5 is proximal to the quencher at the 3C overhang location (Figure 1, D). Thus, the Cy3 and Cy5 are quenched in the initial configuration, and hence emit a low fluorescent signal (Cy3 dark green and Cy5 dark red in Figure 1, E). The addition of the target DNA strands causes the QO strands to be removed via toehold mediated strand displacement^[42]^ (Figure 1, D). After target A (DNA target corresponding to Cy3 channel) displaces the QO, the Cy3 molecule emits higher fluorescent signal as is shown with the dashed green spectra in Figure 1, E. The Cy3 fluorescent signal of the CSP folded without the quenching oligo was also shown as a control to display the maximum fluorescent signal (light green spectra in Figure 1, E). The design principles are the same for the Cy5-labeled oligo and 3C overhang. The Cy5 fluorescent signal is low before the addition of target B (DNA target corresponding to Cy5 channel) as shown with solid dark red spectra in Figure 1, E. Incubation of target B with the CSP causes strand displacement and an increase in the Cy5 fluorescent signal (dashed red spectra in Figure 1, E). The light red spectra in Figure 1, E shows the Cy5 signal from the CSP folded without QO, as the control showing essentially all quencher strands are displaced.

### 2.2: CSP detects DNA targets on the cell membrane

To detect DNA targets on the membrane of living cells, the CSP was incorporated onto the extracellular side of cell membranes using cholesterol-conjugated-oligonucleotides based on a previously published method^[20]^ (Figure 2, A). The functionalized cells with CSP were immobilized on a glass chamber and either imaged directly using epifluorescence microscopy or imaged after introducing 1uM target nucleic acid strands into solution and incubating for 15 min at 37°C. Representative fluorescent images in Figure 2, B show: I) cells functionalized with non-QO labeled CSP, II) cells functionalized with QO labeled CSP, III) cells with QO labeled CSP after incubation with target A, IV) cells with QO labeled CSP after incubation with target B, V) cells with QO labeled CSP after incubation with both DNA targets, and VI) cells functionalized with non-QO labeled CSP after incubation with both DNA targets.

**Figure 2:**
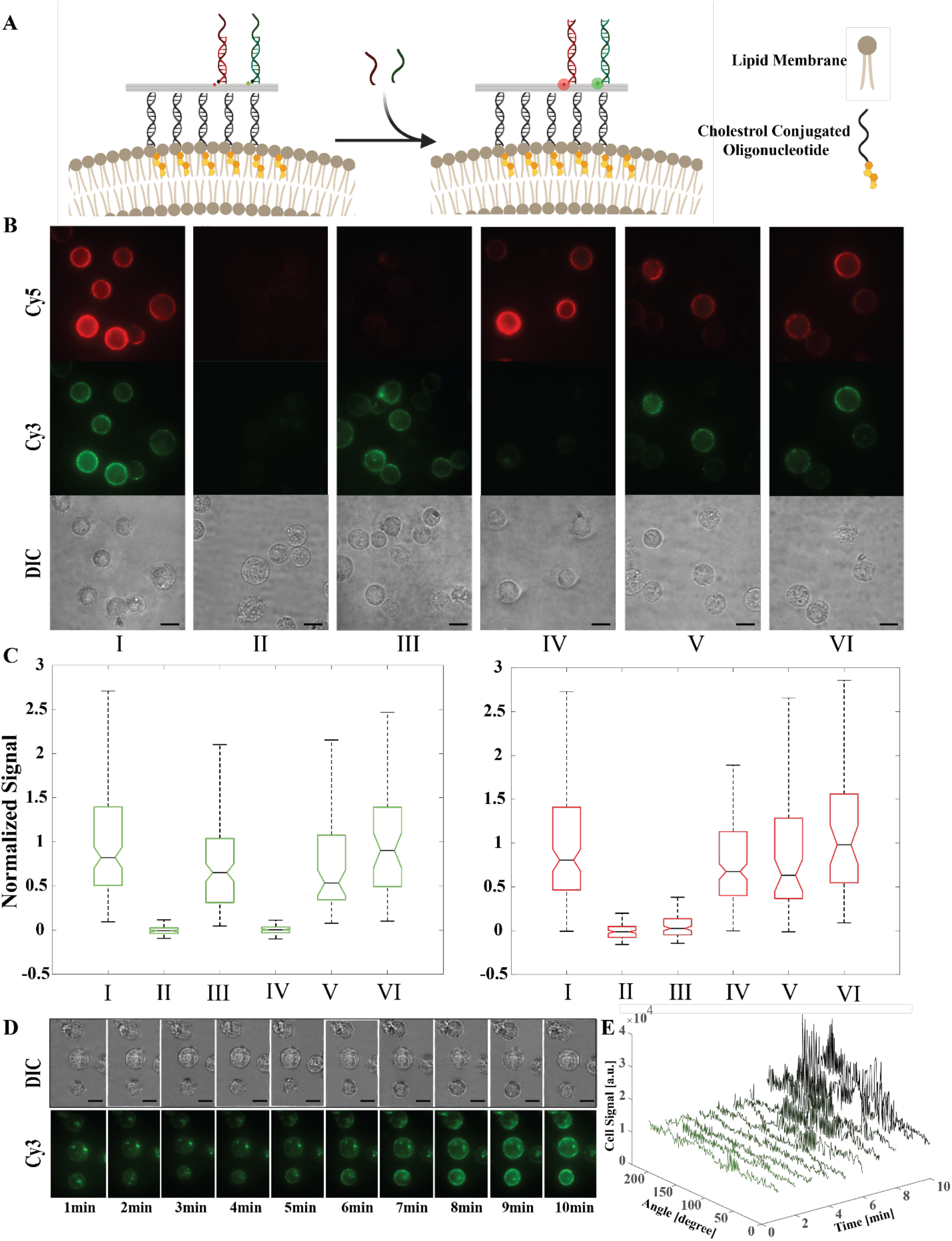
Detection of ssDNA targets on CH12-LX suspension cells. A) Schematics of the detection of ssDNA targets on the cell membrane. CSP is incorporated into the cell membrane. The addition of DNA targets, results in the displacement of QO and high fluorescent signal. B) Fluorescent and DIC images representing controls and steps taken to detect target oligos on the cell membrane. I) CH12-LX cells functionalized with non-QO labeled CSP (control). II) CH12-LX cells functionalized with QO labeled CSP. III) sample in (II) after the addition of target A. IV) Sample in (II) after the addition of target B. V) Sample in (II) after the addition of both targets. VI) Sample in (I) after the addition of both targets (control). Scale bars, 100um. C) The mean fluorescence intensity attributed to the CSP bound to the surface of 100–300 single cells was quantified based on three independent experiments for each condition. The data are expressed as cell’s mean fluorescent intensity normalized relative to the average of the mean fluorescence intensity quantified for conditions I and II (red: Cy5 signal, green: Cy3 signal). D) Detection of target A on the membrane of CH12-LX cells in real-time. Left: DIC and fluorescent images show the increase of Cy3 signal during 10 min. Right: The fluorescence intensity attributed to the CSP bound to the surface of the middle cell was quantified for each time point around the membrane (Scale bars, 100um).

The fluorescence intensity attributed to CSP on the surface of cells was measured using a custom MATLAB code and parameterized in terms of the mean fluorescence intensity around the perimeter of individual cells^[20]^. The mean fluorescence intensity for individual cells was normalized with respect to the overall average of the mean fluorescence intensity under conditions I (i.e., cells functionalized with non-QO labeled CSP) and II (i.e., cells functionalized with QO labeled CSP) as the maximum and minimum, respectively. Based on the results in Figure 2, C the cells’ mean fluorescence intensity from the CSP increased robustly after the addition of the respective DNA targets A and B compared to condition II (cells functionalized with QO labeled CSP). The overall average of the mean fluorescence intensity of the cells incubated with target A (condition III, Cy3 plot in green) has a similar average intensity to cells functionalized with non-QO labeled CSP (condition I, Cy3 plot in green). Similarly, the average from the cells functionalized with QO labeled CSP and incubated with target B (condition IV, Cy5 plot in red) is as high as the average in cells functionalized with non-QO labeled CSP (condition I, Cy5 plot in red). Importantly, comparing cells with QO labeled CSP after incubation with targets A or B (conditions III and IV) to the minimal signal (i.e., cells functionalized with QO labeled CSP, condition II) shows that the addition of each DNA target only affects the corresponding fluorescent signal and does not affect the fluorescent intensity in the other channel. In addition, exposing cells with QO labeled CSP to both targets (condition V) leads to robust fluorescence increases in both fluorescence channels, illustrating both specific and multiplexed sensing of molecular targets.

To demonstrate CSP as a tool to monitor cell-biomolecule interactions in real time, we demonstrated the detection of target A on the cell membrane over a time span of 10 min. Figure 2, D shows brightfield and fluorescent images of 3 single cells, which were initially functionalized with QO labeled CSP. At time t=0 min, target A was added to the imaging dish at a final concentration of 1uM, and images were taken every minute to monitor the temporal increase in CSP fluorescence intensity due to detection of target A. We quantified the fluorescent intensity around the membrane of the middle cell at each time point (Figure 2, E). The result shows up to 4-fold increase of the Cy3 intensity at some angles, which highlights the ability to resolve interactions with sub-cellular resolution around the cell surface. The nonuniform fluorescent intensity increase on the cell periphery could be a result of the inhomogeneous distributions of CSP around the cell membrane. However, the non-QO labeled CSP conditions show a relatively uniform density of structures around the periphery of cells, suggesting the inhomogeneous reporter fluorescence may be due to non-uniform binding of targets.

### 2.3: CSP detects cell membrane interactions in 3-D ECM model

Interactions between cells and their microenvironment are highly affected by ECM biophysical and biochemical properties. Design and fabrication of integrated microfluidic devices with localized 3-D ECM compartments are advancing the study complicated living systems^[43,44]^, hence, aiding understanding of detailed biological mechanisms. These devices enable controlling biophysical properties of the microenvironment such as collagen density, while enabling measurements with subcellular spatial resolution in real time. Here, we used a tissue model to establish the successful incorporation of functionalized cells into the collagen matrix to investigate the cell-target molecule interaction in a 3D microenvironment that is more representative of native biological conditions.

To determine how target biomolecules interact with functionalized cells in the 3-D collagen matrix, we developed a microfluidic platform that features fluid flowing through two adjacent channels, which we call side channels, on either side of a central channel that contains a 3-D collagen extracellular matrix (ECM). We refer to this microfluidic system as an ECM Probing Chip (EPC). The EPC has two side channels (50 μm in height) that are spaced 1 mm apart, each with its own individual inlet and outlet. The individual inlets and outlets in the side channels (Figure 3, A - purple channels) allow for the control of flow through the side channels and across the middle ECM channel, which contains a collagen matrix that can be seeded with live cells. Along the device, there are six apertures (100 μm in width and 50 μm apart) that allow for components introduced into a side channel (e.g., target strands) to flow through the 3-D collagen matrix (Figure 3, A - green channel). In these experiments, dendritic cells were functionalized and mixed with collagen I and were seeded into the middle ECM channel of the EPC and incubated for 30 min at 37°C to ensure collagen polymerization prior to running detection experiments.

**Figure 3:**
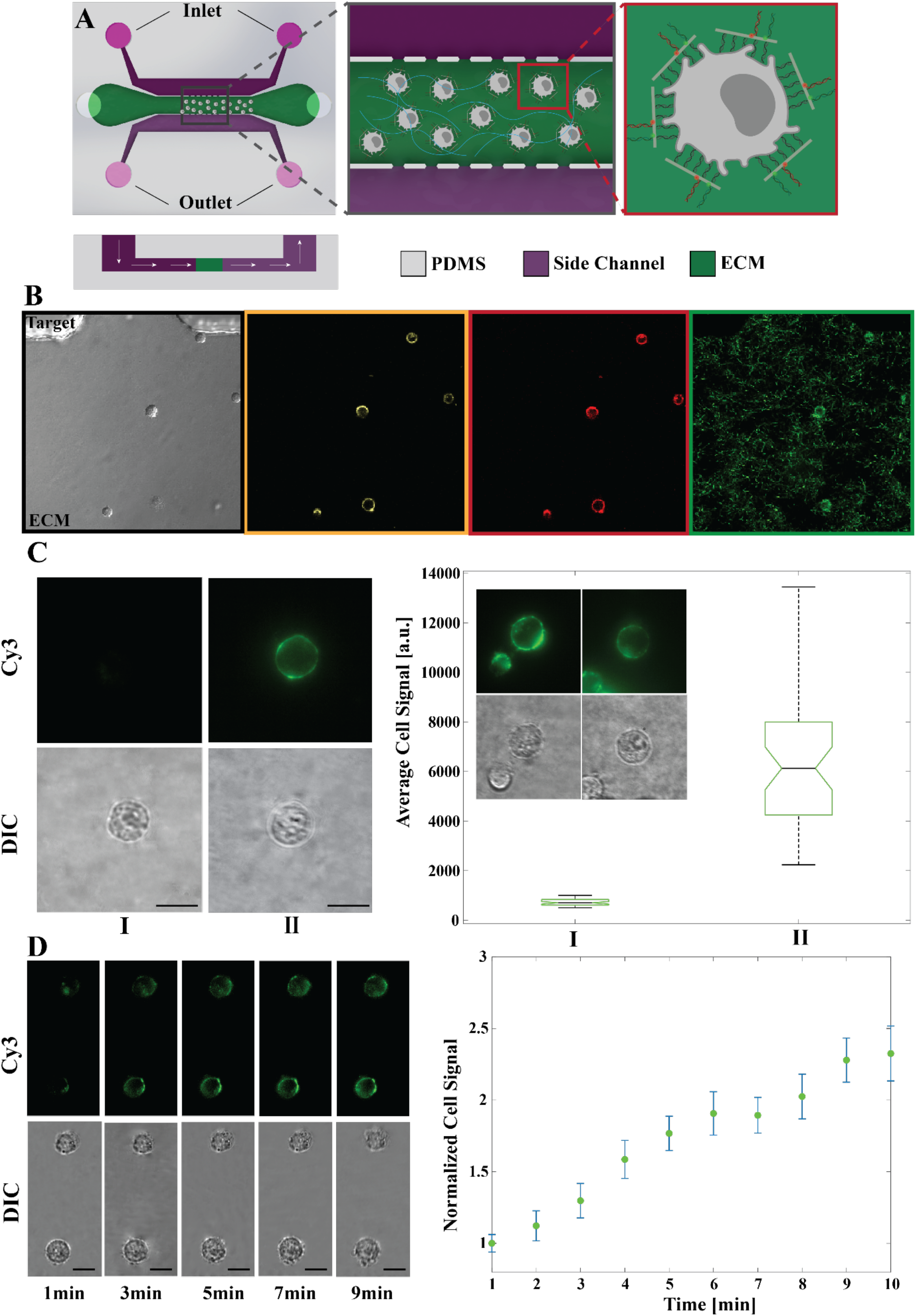
Detection of ssDNA targets on DCs in a 3-D collagen matrix using the EPC. A) Schematic images of the microfluidic platform used for the detection. Left) Top view of the channel featuring localized region of mixture of cells and the collagen gel (green) and the channels that will be used to apply the flow of targets (purple). Each side channel has independent input and outlet ports, allowing the control of flow in both channels. Middle) Close-up view of boxed area in Left showing apertures that allow the connection of the target channels with cell channel. Right) Close-up view of boxed area in middle showing cells functionalized with CSP in collagen gel seeded into the middle channel of the EPC. B) Confocal images representing cells functionalized with CSP in collagen gel seeded into the middle channel of the EPC. Left) Bright field image showing cells seeded into the middle channel and one aperture connecting the middle channel to the top channel. Using confocal microscopy, the formation of collagen fibers (green), successful binding of CSP to the cells (red and yellow channels) and incorporation of cells into the collagen matrix is shown. C) Fluorescent and DIC images representing the detection of target A on the cell membrane in the collagen matrix. I: DC before the addition of target A. II: DC after the flow of target A into the collagen channel. Box plot represents the mean fluorescence intensity of ~50 cells (for each condition) in the collagen matrix, before and after the flow of target in 5 different EPC. The individual cells are examples of DCs functionalized with QO after the addition of target A (II). D) Detection of target A on the surface of DC in the collagen matrix in real-time. DIC and fluorescent images show the increase of Cy3 signal over 10 min (left). The average of Cy3 signal on the membrane of seven cells, seeded into three different EPC over 10 minutes. The fluorescent signal is normalized based on the average signal of all the cells at time 0 and 10 min, and the rate of photobleaching.

First, we showed with fluorescence and confocal microscopy the successful formation of collagen fibers and the stability of functionalized cells incorporated into collagen matrices. Figure 3, B shows (left to right) brightfield, Cy3, Cy5 and reflectance images of DC incorporated into the middle channel, located close to an aperture. Yellow and Red images demonstrate the stability of CSP on the membrane of the DCs at 2 hours after the preparation of the samples and the reflectance image shows the collagen matrix structure.

In order to establish the functionality of CSP on the membrane of DC, we repeated the two-channel detection experiment with DCs. Figure S4 shows the successful multiplexed detection of targets A and B on the membrane of DCs deposited in an imaging dish. Next, we tested the detection of target A on the DCs seeded within the ECM in the EPC. DCs functionalized with QO labeled CSP were seeded into the middle channel of 5 EPC devices, and DIC and fluorescent images were taken from ~50 individual cells (condition I in Figure 3, C). Lastly, 1uM of target A was introduced into the inlet channel (target channel) and 1xPBS was added to outlet channel. By applying a droplet of DNA target on top of both reservoirs of the inlet channel, a pressure difference was applied across the collagen channel, causing flow of the DNA target through the collagen. Near 50 individual cells were again imaged after 15 min (condition II in Figure 3, C) and the mean fluorescence intensity was measured. The box plot in Figure 3, C shows the average of DCs mean fluorescence intensity increased by approximately 6 times after the addition of the target DNA.

We further demonstrated the capability of our platform to study the interaction of biomolecules with the cell membrane in situ in real time. The EPC was seeded with DCs functionalized with QO labeled CSP and was mounted on the imaging stage. Target DNA and 1xPBS were introduced into the inlet and outlet channels, respectively. Then, two droplets of DNA target were applied on top of the inlet reservoirs to help drive the flow through the middle channel and individual cells were imaged every minute over 10 minutes. Figure 3, D shows DIC and fluorescence images of two CSP functionalized DCs imaged over time. The intensity of the Cy3 signal increased significantly over 10 minutes after introduction of the target DNA. The mean fluorescence intensity of seven individual cells seeded in 3 EPCs at each time point was measured and the plot in Figure 3, D shows its average relative to the average signal at time zero over the timespan of the experiment. We normalized individual cell’s mean Cy3 signal with respect to the average Cy3 signal at time 0. To account for photobleaching, we also normalized the mean fluorescence intensity of each cell at each timepoint to the average of cell fluorescence intensity for the case where no target was added to cells with non-QO labeled CSP (Figure S5). The Cy3 reporter signal increases steadily, more than doubling over 10 minutes. This experiment was repeated, and cells were imaged for 50 minutes, with 5-minute intervals (to prevent significant photobleaching), and results are shown in Figure S6. These experiments revealed the signal increased mainly over the first 10 min of the experiment and saturated by approximately 20 min. These results demonstrate the ability to study dynamic cell-biomolecule interactions over time, with higher time resolution over short times, or over prolonged periods of time with lower time resolution.

## 3. Conclusion

The ability to monitor the interactions of cells with the instructive cues originating from the local molecular and extracellular environment is essential for understanding the determinants of cell fate and function,^[45]^ for example, regulating cellular spatiotemporal organization^[46,47]^, processes such as neurotransmission^[48]^, wound healing, and inflammation^[49]^, and progression of diseases including metabolic disorders, autoimmune diseases, and cancer^[50–53]^. Here, we established a method to functionalize living cells with DNA origami sensors to spatiotemporally monitor interactions of cells with biomolecules in the local ECM microenvironment using our microfluidic 3-D tissue model device. We demonstrated the multiplexed sensing on the surface of CH12-LX B cells and demonstrated the capability of our method to monitor subcellular interactions of MutuDC 1949 cells in the 3D ECM. Our results established the integration of DNA origami nanodevices, cell membrane engineering, and microfluidic tissue model systems as a powerful approach to probe biological interactions in physiologically relevant 3-D environments with high spatial (i.e. micron-scale) and temporal (i.e. minute scale) resolution. The time resolution could easily be improved to the second scale to study faster interactions; however, this could require additional strategies to inhibit photobleaching ^[54,55]^.

Here, we focused on nucleic acid targets as a proof-of-concept to enable the detection of relevant targets such as miRNA and circulating tumor DNA (ctDNA) that are present in extracellular circulation and ECM and can play an important role in activation of immune cells^[56,57]^. This approach could be expanded to study cell response to a variety of molecules. For example, the incorporation of aptamers could enable the detection of targets growth factors or cytokines^[58]^ and the measurement of local environmental factors such as pH^[59]^. Other DNA constructs have been demonstrated as tools to measure forces^[19,21,60,61]^ or ion concentrations^[62]^. The programmability and stability of the platform in a wide range of solution conditions allows for the incorporation and functionality of many DNA aptamers^[8]^. Thus, it is possible to look at the underlying mechanisms of cooperative action between multiple types of biomolecules or biomolecules and other local factors (pH, forces, ions).

In this study, we multiplexed two organic fluorophores to label our DNA nanostructures. Common fluorescence imaging systems could multiplex 3-4 channels. Further multiplexing could be achieved using novel fluorescent imaging techniques such as metafluorophores^[63]^, frequency multiplexed DNA-PAINT^[64]^, or fluorescent nanoparticles with controllable spectral properties^[65]^ that could expand the distinct number of channels, and minimize cross-talk and photobleaching. For examples, luminescent quantum dots (QDs) have been proven as a promising alternative to traditional organic dyes in various fluorescence-based applications such as multicolor live cell imaging over a week^[66]^, and QDs have been used for Förster Resonance Energy Transfer (FRET) based studies of biomolecular interactions^[67]^.

We successfully established a reliable method to study subcellular interactions in a 3-D tissue model using a microfluidic device. Microfluidic devices are enabling novel approaches for probing important biological questions in cellular microenvironments with precise control over biophysical and biochemical parameters ^[68]^. Moreover, cutting-edge microfluidic models allow for recapitulating native cell-ECM structures in microfluidic that orchestrate physiological processes such as angiogenesis, vessel branching, and tissue morphogenesis. Prior work has utilized microfluidic devices to actuate DNA origami nanodevices inside cell-sized microfluidic compartments^[69]^. Here we establish the integration of DNA nanodevices, cell membrane engineering, and microfluidic tissue model systems to enable a unique approach to study subcellular biological phenomena in a 3D ECM. Combining the programmability of DNA origami with versatile designs of microfluidic chips allows for the investigation of the subcellular interactions under controlled biomolecular transport and fluid mechanical stimuli with unprecedented spatial and temporal resolution.

## 4. Experimental Section

### 4.1: Design and Fabrication of the CSP

The CSP was designed using the software caDNAno^[40]^ and fabricated using protocols developed by Castro et al^[70]^. CSP is a 65nm×30nm×6nm platform, with 42 potential overhangs (Figure 1). Staple sequences were specified in caDNAno and ordered from a commercial vendor (Integrated DNA Technologies, Coralville, IA). The CSP was folded using p7249 from the M13mp18 genome with a total of 186 ssDNA staples. To fold the CSP with 2 DNA detection modules on locations C3 and C4, staple strands with an end directly adjacent to the overhang locations (Green oligos in Figure S1, and Table S1) oligos on these two locations were replaced with one DNA oligo labeled with Cy5 and one DNA oligo labeled with Cy3 fluorophore (Table S2). Moreover, CSP was folded with 30 overhangs on the membrane facing side (bottom overhangs, purple oligos in Figure S1 and Table S1) and 2 overhangs on the opposite side (top overhangs, Table S2) on locations C3 and C4. Briefly, purified scaffold at 20 nM concentration was combined with 5-fold molar excess of staples (each staple at 100 nM), and 10-fold molar excess of QO (if needed, in 200nM each) in folding buffer (1xFOB: 5 mM Tris, 5 mM NaCl, 1 mM EDTA, supplemented with 18mM MgCl_2_).

In order to find the optimal salt concentration needed for structural folding, initially folding reactions with different salt concentrations (10, 12, 14, 16, 18, 20, 22, 24 mM MgCl2) were prepared. Then the self-assembly reaction was done by rapidly heating the reactions to 65oC followed by slow cooling to 4°C over 2.5 days in a thermal cycler (Bio Rad, Hercules, CA). The folding reaction was brought to 65°C and stepped from 65°C to 24°C in 1°C steps at varying time increments. From 65°C to 62°C, the temperature was held for 1 hour; from 61 °C to 59°C, the temperature was held for 2 hours; from 58°C to 46°C, the temperature was held for 3 hours; from 45°C to 40°C, the temperature was held for 1 hour; from 39°C to 24°C, the temperature was held for thirty minutes. After being held at 24°C for 1 hour, the temperature was lowered to 4°C. Folding reaction products were subjected to agarose gel electrophoresis^[15]^ in a 2% agarose (Life Technologies) gel (0.5x TBE) in the presence of 11 mM MgCl2 and 1 μM ethidium bromide (EtBr) and based on the results, 18Mm MgCl_2_ was chosen as the optimized concentration for folding the CSP (Figure S1).

Next, using folding reactions at 18Mm MgCl_2_, the optimal isothermal annealing temperature was found. The folding mixtures were subjected to thermal annealing by rapid heating to 65 °C followed by isothermal annealing for 4 hours in different temperatures (60, 58.4, 56.0, 52.5, 47.8, 44.4, 41.7, 40) a rapid cooling to 4 °C. Folding reaction products were subjected to agarose gel electrophoresis and the results revealed the successful folding of CSP over a wide range of 42 to 59°C (Figure S1). 52°C was chosen for subsequent folding with isothermal annealing. Therefore, to fold CSP, the folding mixtures at 18mM MgCl_2_ were subjected to thermal annealing by rapid heating to 65 °C followed by 4 hours in 52 degrees and a rapid cooling to 4 °C.

### 4.2: DNA Nanostructure Purification

Folded DNA nanostructures were purified by mixing folding reaction products with equivolume of 15% PEG 8000 (Sigma-Aldrich, St Louis, MO) supplemented with 500 mM NaCl and centrifugation for 30 minutes at 16000 rcg to remove excess staple strands (protocol modified from^[71]^). After removing the supernatant, purified structures were resuspended in 1xFOB supplemented with 10 mM MgCl_2_ (storage buffer). To fully remove the to remove excess staple strands, the PEG purification procedure was repeated two times.

### 4.3: Transmission Electron Microscopy (TEM)

TEM grids were prepared as described in Castro *et al*.^[70]^. Briefly, 4μL of ~1 nM purified DNA nanostructure was deposited onto a copper TEM grid coated with carbon and formvar (Electron Microscopy Sciences, Hartfield, PA) and incubated for 4 minutes at room temperature. The sample was removed by gently touching filter paper to the edge of the grid and 10μL of 2% uranyl formate negative stain was applied to the grid and immediately removed with filter paper as a washing step. Then, 20μL of 2% uranyl formate was immediately added, incubated for 40 seconds, and finally removed with filter paper. The grid was allowed to dry for at least 30 minutes before being visualized on a Tecnai G2 BioTWIN transmission electron microscope (FEI, Hillsboro, OR) in the Ohio State University Campus Microscopy & Imaging Facility (CMIF).

### 4.4: Functionality of CSP in detection of DNA targets

To evaluate CSP strand displacement, structures (folded with QO) at 5 nM concentration were incubated with 1uM final concentration of each DNA target (A or B) at 37 °C for 15 min followed by a fluorescent measurement on fluorometer (FluoroMax-4, Horiba, Japan). The samples were excited with 510nm and 610nm lasers for target A and target B respectively and the emission was measured from 530nm to 700nm for target A, and 630nm to 700nm for target B. The fluorescent intensity of CSP folded with or without QO in the same concentration (5nM) was also tested on the fluorometer to demonstrate the maximum and minimum fluorescent signal.

### 4.5: Stability of the CSP in cell culture condition

To confirm CSP structural integrity under cell culture conditions, folded structures were purified using PEG centrifugal purification process and resuspended in different cell culture media and were incubated for 4 hours at 37 °C. Purified structures were incubated in CH12-LX clear cell culture media (RPMI 1640 without L-glutamine (Corning)) supplemented with 1 mM MgCl_2_, RPMI supplemented with 2% heat-inactivated Fetal Bovine Serum (FBS) (Atlas Biologicals) and 1mM MgCl_2_, RPMI supplemented with 8% FBS and 1mM MgCl_2_, 1 x Phosphate Buffered Saline (PBS) without MgCl_2_/CaCl_2_ (Corning, Cat# 21-031-CV), 1 x PBS supplemented with 2% FBS and 1mM MgCl_2_, and 1 x PBS supplemented with 8% FBS and 1mM MgCl2 following PEG purification. Agarose gel electrophoresis assay in a 2% agarose gel in the presence of 11 mM MgCl2 and 1 μM ethidium bromide followed by visualization of the excised gel bands on TEM was used to confirm the stability of the structures in the cell culture media (Figure 1 and S3).

### 4.6: Cell culture

CH12-LX is a murine B-cell lymphoma obtained from Dr. Gail Bishop^[72]^. In order to culture the CH12-LX B cells, the RPMI 1640 without L-glutamine (Corning) was supplemented with 10% heat-inactivated FBS (Atlas Biologicals), 1% Penicillin-Streptomycin-Glutamine (100X) (Thermo-Fisher), 1% Sodium Pyruvate (100mM) (Life Technologies), 1% HEPES (1M) (Life-Technologies), 1% MEM Non-Essential Amino Acids (100X) (Life Technologies). For experiments, CH12-LX cells were washed once with 1x PBS, followed by one wash with the experimental medium (clear RPMI 1640 without L-glutamine (Corning) supplemented with 2% heat-inactivated FBS, and 1mM MgCl_2_). Finally, CH12-LX cells were resuspended in the experimental medium at 4000 cells/uL for incubation with cholesterol-conjugated oligos.

The wild-type MutuDC1940 dendritic cell (DC) line is derived from mouse spleen tissues and is GFP positive due to the GFP reporter in the CD11c:SV40LgT transgene. This cell line was obtained from abm (Richmond BC, Canada) and were maintained in supplemented IMDM in 37°C incubator at 5% CO_2_. To make the complete growth medium, IMDM (1x) + GlutamaxTM (Gibco Ref: 31980-030) was supplemented with 10% heat-inactivated Fetal Bovine Serum (FBS) (Atlas Biologicals), 1% of 7.5% Sodium Bicarbonate Solution (Life Technologies), 50 μM β-mercaptoethanol, 1% HEPES (1M) (Life-Technologies), and 1% Penicillin-Streptomycin-Glutamine (100X) (Thermo-Fisher). For passaging purposes, the cells were washed with 1 x PBS without MgCl_2_/CaCl_2_, followed by their detachment from the culture flask using 1:1 ratio of 1 x PBS and 0.25% Trypsin-EDTA (1X) (Corning). The cells were incubated with PBS-Trypsin mixture for 3-4 minutes at room temperature. The Trypsin was then neutralized using the culture media. For functionalization experiments, cells were washed once with 1 x PBS followed by one wash with the experimental medium (clear RPMI 1640 without L-glutamine (Corning) supplemented with 2% heat-inactivated Fetal Bovine Serum (FBS) (Atlas Biologicals), and 1mM MgCl_2_). Finally, DCs were resuspended in the experimental medium at 4000 cells/uL for incubation with cholesterol-conjugated oligos.

### 4.7: Cell membrane functionalization

Cells were functionalized with CSP using the protocol explained in Akbari *et. al*^[20]^. Briefly, cells resuspended in the experimental medium (clear RPMI 1640 without L-glutamine (Corning) supplemented with 2% heat-inactivated FBS (Atlas Biologicals), and 1mM MgCl_2_) were incubated with 10 μM cholesterol-conjugated oligonucleotide (Table S3) for 5 minutes at 37°C. The cells were then washed once in the experimental media to remove the excess cholesterol-conjugated oligonucleotide. Cell were then incubated with the 60-base bridge oligo at 1 μM for 5 minutes at 37°C, followed by the addition of the 20-base pair fortifier oligos at 1μM, incubation for 5 minutes at 37°C and a wash with experimental media. Finally, the cells were incubated with the nanostructures at 5nM, for 5 minutes at 37°C. Cells were washed a final time to remove excess nanostructures using the experimental media and were resuspended in either 1xPBS or the experimental media for the experiment.

### 4.8: DNA target detection on the membrane of suspension cells

CSP folded with QO and without QO were incorporated into the membrane of two separate subpopulations of cells (CH12-LX or DCs). The subpopulation with QO then were divided into four smaller subpopulations, transferred to an 8-well imaging dish, and incubated with either Cy3-target at 1uM concentration, Cy5-target at 1uM concentration, both DNA targets (A and B) at 1uM final concentration or 2uL ddH_2_O (to maintain the same buffer composition) in 37 °C for 15min. Cells functionalized with CSP devices without QO were divided into two subpopulations and incubated with either 1uM final concentration of each DNA targets A or B or 2uL ddH_2_O on the imaging dish in the same condition. All six conditions were imaged using DIC, Cy3 and Cy5 imaging settings (see supplemental info). For real-time experiments, the Cy3-target was added at 1uM final concentration to the subpopulation of cells functionalized with QO on the microscope stage after capturing the first image at time=0. The DIC and Cy3-fluorescence images were captured every minute. The fluorescent signal on the membrane of each cell was measured and analyzed using a home-built MATLAB code described below.

### 4.9 DNA target detection on the membrane of DCs seeded into the EPC

DCs were functionalized with CSP using the same protocol from ref^[20]^. While preparing the cells, a collagen mixture was prepared according to vendor’s instructions. Briefly, the pH of high concentration rat-tail collagen I stored in acidic solution (Corning Life Sciences) was neutralized to a pH of 7.4 using NaOH in 10 x PBS without MgCl_2_/CaCl_2_ so that the final concentration of the PBS is 1x. The mixture was prepared in a 4 °C ice bath and incubated at the same temperature for 10 min. Finally, functionalized cells were resuspended in 1 x PBS without MgCl_2_/CaCl_2_ and mixed with collagen so that the concentration of collagen and cells are 2 mg/ml, and 7000 (cells/uL), respectively. The final mixture was seeded into the middle channel of the EPC and incubated at 37 °C for 30 min to allow for formation of collagen matrix before imaging. Each EPC was mounted onto imaging stage and approximately 20 cells were imaged using DIC and Cy3-fluorescence settings. DNA target A at 1uM concentration and 1xPBS were added to the EPC inlet and outlet channels respectively, and the EPC was incubated for 15 min at room temperature. Then, 20 cells were imaged using the same settings to study DNA displacement on their membrane. For real-time experiments, the Cy3-target A at 1uM concentration and 1xPBS were added to the EPC inlet and outlet channels on the microscope stage after capturing the first image at time=0. The DIC and Cy3-fluorescence images were captured either every minute or every 5 minutes. The fluorescent signal on the membrane of each cell was measured and analyzed using a home-built MATLAB code.

### 4.10: Cell Periphery Fluorescent Analysis

A home-built MATLAB code previously developed by Akbari *et. al*^[20]^ was used to analyze the fluorescent signal on the membrane of the cells. The code measures the intensity of each pixel around the circumference of each cell and reports the mean fluorescent signal along with the angular distribution of the fluorescent signal. Between 100 and 300 cells were analyzed for each suspension experiments and near 50 cells were analyzed for collagen seeded cell experiments.

## Acknowledgments

This work was supported by the National Institute of Health (grant R01HL141941 awarded to JWS and CEC). The Electron and Confocal microscopy were done at Campus Microscopy & Imaging Facility (CMIF), the Ohio State University. The clean room microfabrication procedure was performed at Nanotech West Laboratory, the Ohio State University. MS and PEB are a recipient of the Pelotonia Fellowship from The Ohio State University Comprehensive Cancer Center.

## References

[1] P. Blanco, A. K. Palucka, V. Pascual, J. Banchereau, Cytokine Growth Factor Rev. 2008, 19, 41.

[2] H. Liang, K. Kidder, Y. Liu, ExRNA 2019, 1, 9.

[3] P. Ranganathan, A. Ngankeu, N. C. Zitzer, P. Leoncini, X. Yu, L. Casadei, K. Challagundla, D. K. Reichenbach, S. Garman, A. S. Ruppert, S. Volinia, J. Hofstetter, Y. A. Efebera, S. M. Devine, B. R. Blazar, M. Fabbri, R. Garzon, J. Immunol. 2017, 198, 2500 LP.

[4] T. H. Kang, C.-P. Mao, Y. S. Kim, T. W. Kim, A. Yang, B. Lam, S.-H. Tseng, E. Farmer, Y.-M. Park, C.-F. Hung, J. Immunother. Cancer 2019, 7, 260.

[5] S. T. Laughlin, J. M. Baskin, S. L. Amacher, C. R. Bertozzi, Science (80-.). 2008, 320, 664 LP.

[6] A. J. R. Amaral, G. Pasparakis, Acta Biomater. 2019, 90, 21.

[7] B. N. G. Giepmans, S. R. Adams, M. H. Ellisman, R. Y. Tsien, Science (80-.). 2006, 312, 217 LP.

[8] W. Zhao, S. Schafer, J. Choi, Y. J. Yamanaka, M. L. Lombardi, S. Bose, A. L. Carlson, J. A. Phillips, W. Teo, I. A. Droujinine, C. H. Cui, R. K. Jain, J. Lammerding, J. C. Love, C. P. Lin, D. Sarkar, R. Karnik, J. M. Karp, Nat. Nanotechnol. 2011, 6, 524.

[9] X. K. Lun, B. Bodenmiller, Mol. Cell. Proteomics 2020, 19, 744.

[10] J. M. Muncie, V. M. Weaver, Curr. Top. Dev. Biol. 2018, 130, 1.

[11] P. E. Beshay, M. G. Cortes-Medina, M. M. Menyhert, J. W. Song, Adv. NanoBiomed Res. 2022, 2, 2100056.

[12] N. C. Seeman, J. Theor. Biol. 1982, 99, 237.

[13] S. Huo, H. Li, A. J. Boersma, A. Herrmann, Adv. Sci. 2019, 6, 1900043.

[14] P. W. K. Rothemund, Nature 2006, 440, 297.

[15] C. E. Castro, F. Kilchherr, D.-N. Kim, E. L. Shiao, T. Wauer, P. Wortmann, M. Bathe, H. Dietz, Nat. Methods 2011, 8, 221.

[16] S. M. Douglas, H. Dietz, T. Liedl, B. Högberg, F. Graf, W. M. Shih, Nature 2009, 459, 414.

[17] A. E. Marras, L. Zhou, H.-J. Su, C. E. Castro, Proc. Natl. Acad. Sci. U. S. A. 2015, 112, 713.

[18] L. Zhou, A. E. Marras, H. J. Su, C. E. Castro, ACS Nano 2014, 8, 27.

[19] M. W. Hudoba, Y. Luo, A. Zacharias, M. G. Poirier, C. E. Castro, ACS Nano 2017, 11, 6566.

[20] E. Akbari, M. Y. Mollica, C. R. Lucas, S. M. Bushman, R. A. Patton, M. Shahhosseini, J. W. Song, C. E. Castro, Adv. Mater. 2017, 29, DOI 10.1002/adma.201703632.

[21] J. V. Le, Y. Luo, M. A. Darcy, C. R. Lucas, M. F. Goodwin, M. G. Poirier, C. E. Castro, ACS Nano 2016, 10, 7073.

[22] B. Ding, Z. Deng, H. Yan, S. Cabrini, R. N. Zuckermann, J. Bokor, J. Am. Chem. Soc. 2010, 132, 3248.

[23] R. Chhabra, J. Sharma, Y. Ke, Y. Liu, S. Rinker, S. Lindsay, H. Yan, J. Am. Chem. Soc. 2007, 129, 10304.

[24] H. T. Maune, S. Han, R. D. Barish, M. Bockrath, W. A. G. III, P. W. K. Rothemund, E. Winfree, Nat. Nanotechnol. 2010, 5, 61.

[25] H. Bui, C. Onodera, C. Kidwell, Y. Tan, E. Graugnard, W. Kuang, J. Lee, W. B. Knowlton, B. Yurke, W. L. Hughes, Nano Lett. 2010, 10, 3367.

[26] S. M. Douglas, I. Bachelet, G. M. Church, Science 2012, 335, 831.

[27] Q. Zhang, Q. Jiang, N. Li, L. Dai, Q. Liu, L. Song, J. Wang, Y. Li, J. Tian, B. Ding, Y. Du, ACS Nano 2014, 8, 6633.

[28] Y.-X. Zhao, A. Shaw, X. Zeng, E. Benson, A. M. Nyström, B. Högberg, ACS Nano 2012, 6, 8684.

[29] P. D. Halley, C. R. Lucas, E. M. McWilliams, M. J. Webber, R. A. Patton, C. Kural, D. M. Lucas, J. C. Byrd, C. E. Castro, Small 2016, 12, 308.

[30] M. Langecker, V. Arnaut, T. G. Martin, J. List, S. Renner, M. Mayer, H. Dietz, F. C. Simmel, Science (80-.). 2012, 338, 932.

[31] J. R. Burns, A. Seifert, N. Fertig, S. Howorka, Nat. Nanotechnol. 2016, 11, 152.

[32] R. Jungmann, M. S. Avendaño, J. B. Woehrstein, M. Dai, W. M. Shih, P. Yin, Nat. Methods 2014, 11, 313.

[33] D. Wang, Y. Fu, J. Yan, B. Zhao, B. Dai, J. Chao, H. Liu, D. He, Y. Zhang, C. Fan, S. Song, Anal. Chem. 2014, 86, 1932.

[34] P. C. Nickels, B. Wünsch, P. Holzmeister, W. Bae, L. M. Kneer, D. Grohmann, P. Tinnefeld, T. Liedl, Science (80-.). 2016, 354, 305.

[35] J. Li, K. Xun, K. Pei, X. Liu, X. Peng, Y. Du, L. Qiu, W. Tan, J. Am. Chem. Soc. 2019, 141, 18013.

[36] H. G. Franquelim, A. Khmelinskaia, J.-P. Sobczak, H. Dietz, P. Schwille, Nat. Commun. 2018, 9, 811.

[37] R. Rubio-Sánchez, S. E. Barker, M. Walczak, P. Cicuta, L. Di Michele, Nano Lett. 2021, 21, 2800.

[38] V. J. Schüller, S. Heidegger, N. Sandholzer, P. C. Nickels, N. A. Suhartha, S. Endres, C. Bourquin, T. Liedl, ACS Nano 2011, 5, 9696.

[39] Y. Wang, E. Benson, F. Fördős, M. Lolaico, I. Baars, T. Fang, A. I. Teixeira, B. Högberg, Adv. Mater. 2021, 33, 2008457.

[40] S. M. Douglas, A. H. Marblestone, S. Teerapittayanon, A. Vazquez, G. M. Church, W. M. Shih, Nucleic Acids Res. 2009, 37, 5001.

[41] J.-P. J. Sobczak, T. G. Martin, T. Gerling, H. Dietz, Science 2012, 338, 1458.

[42] B. Yurke, A. J. Turberfield, A. P. Mills, F. C. Simmel, J. L. Neumann, Nature 2000, 406, 605.

[43] A. Avendano, M. Cortes-Medina, J. W. Song, Front. Bioeng. Biotechnol. 2019, 7, 6.

[44] C. P. Huang, J. Lu, H. Seon, A. P. Lee, L. A. Flanagan, H.-Y. Kim, A. J. Putnam, N. L. Jeon, Lab Chip 2009, 9, 1740.

[45] L. Ferreira, J. M. Karp, L. Nobre, R. Langer, Cell Stem Cell 2008, 3, 136.

[46] M. E. Birnbaum, J. L. Mendoza, D. K. Sethi, S. Dong, J. Glanville, J. Dobbins, E. Özkan, M. M. Davis, K. W. Wucherpfennig, K. C. Garcia, Cell 2014, 157, 1073.

[47] T. Suzuki, Cell. Mol. Life Sci. 2013, 70, 631.

[48] Alcohol Health Res. World 1997, 21, 107.

[49] S. A. Eming, T. Krieg, J. M. Davidson, J. Invest. Dermatol. 2007, 127, 514.

[50] R. Mayor, C. Carmona-Fontaine, Trends Cell Biol. 2010, 20, 319.

[51] M. M. Burdick, O. J. T. McCarty, S. Jadhav, K. Konstantopoulos, IEEE Eng. Med. Biol. Mag. 2001, 20, 86.

[52] S. Sakaguchi, T. Yamaguchi, T. Nomura, M. Ono, Cell 2008, 133, 775.

[53] L. Lu, W. A. Walker, Am. J. Clin. Nutr. 2001, 73, 1124S.

[54] R. A. Hoebe, C. H. Van Oven, T. W. J. Gadella, P. B. Dhonukshe, C. J. F. Van Noorden, E. M. M. Manders, Nat. Biotechnol. 2007, 25, 249.

[55] A. M. Bogdanov, E. I. Kudryavtseva, K. A. Lukyanov, PLoS One 2012, 7, DOI 10.1371/JOURNAL.PONE.0053004.

[56] J. Raisch, A. Darfeuille-Michaud, H. T. T. Nguyen, World J. Gastroenterol. 2013, 19, 2985.

[57] K. Leone, C. Poggiana, R. Zamarchi, Diagnostics 2018, 8, DOI 10.3390/diagnostics8030059.

[58] M. Kimoto, R. Yamashige, K. Matsunaga, S. Yokoyama, I. Hirao, Nat. Biotechnol. 2013, 31, 453.

[59] S. Modi, S. M. G., D. Goswami, G. D. Gupta, S. Mayor, Y. Krishnan, Nat. Nanotechnol. 2009, 4, 325.

[60] R. Glazier, J. M. Brockman, E. Bartle, A. L. Mattheyses, O. Destaing, K. Salaita, Nat. Commun. 2019 101 2019, 10, 1.

[61] M. Iwaki, S. F. Wickham, K. Ikezaki, T. Yanagida, W. M. Shih, Nat. Commun. 2016 71 2016, 7, 1.

[62] S. Saha, V. Prakash, S. Halder, K. Chakraborty, Y. Krishnan, Nat. Nanotechnol. 2015 107 2015, 10, 645.

[63] J. B. Woehrstein, M. T. Strauss, L. L. Ong, B. Wei, D. Y. Zhang, R. Jungmann, P. Yin, Sci. Adv. 2017, 3, DOI 10.1126/SCIADV.1602128.

[64] P. A. Gómez-García, E. T. Garbacik, J. J. Otterstrom, M. F. Garcia-Parajo, M. Lakadamyali, Proc. Natl. Acad. Sci. 2018, 115, 12991 LP.

[65] F. Boschi, F. de Sanctis, Eur. J. Histochem. 2017, 61, DOI 10.4081/EJH.2017.2830.

[66] J. K. Jaiswal, H. Mattoussi, J. M. Mauro, S. M. Simon, Nat. Biotechnol. 2003, 21, 47.

[67] M. Cardoso Dos Santos, W. R. Algar, I. L. Medintz, N. Hildebrandt, TrAC Trends Anal. Chem. 2020, 125, 115819.

[68] E. W. K. Young, D. J. Beebe, Chem. Soc. Rev. 2010, 39, 1036.

[69] K. Göpfrich, M. J. Urban, C. Frey, I. Platzman, J. P. Spatz, N. Liu, Nano Lett. 2020, 20, 1571.

[70] C. E. Castro, F. Kilchherr, D.-N. Kim, E. L. Shiao, T. Wauer, P. Wortmann, M. Bathe, H. Dietz, Nat. Methods 2011, 8, 221.

[71] E. Stahl, T. G. Martin, F. Praetorius, H. Dietz, Angew. Chemie 2014, 126, 12949.

[72] G. Haughton, L. W. Arnold, G. A. Bishop, T. J. Mercolino, Immunol. Rev. 1986, 93, 35.

