## Supplemental Information for "Multiplexed Detection of Molecular Interactions with DNA Origami Engineered Cells in 3-D Collagen Matrices"

The Ohio State University

E328 Scott Laboratory, 201 W. 19th Ave, Columbus, OH 43210

The Ohio State University

E406 Scott Laboratory, 201 W. 19th Ave, Columbus, OH 43210.

**Methods:**

**Total Internal Reflection (TIRF) Microscopy**

Live cell fluorescent imaging was performed on a Nikon TiE (Belmont, CA). An ultra-thin 8-well imaging plate (LAB-TEK) was used to image the suspension samples. Fluorescent excitation was provided by 561 nm (100 mW source) or 640 nm (50 mW source) lasers for Cy3 and Cy5 channels respectively. Images were acquired using an ANDOR EMCCD camera at 160 nm/px resolution. The background noises on the DIC images were removed and the contrast and brightness were enhanced using Adobe Photoshop.

**Confocal Microscopy**

Confocal images were recorded using a laser scanning confocal microscopy (Nikon A1R) in the Ohio State University Campus Microscopy & Imaging Facility (CMIF). A 40x, 1.3 NA oil immersion objective was used for imaging the microfluidic devices. Reflectance images of collagen fibers were acquired by exciting samples using a 487 nm laser, then collecting the emitted signal at 450 nm. CSP localization images on cells were acquired by exciting fluorescence samples by 561 nm and 639 nm lasers, respectively. Image acquisition was performed using an EMCCD Hamamatsu camera at 1024*1024 resolution.

**Fabrication of the microfluidic platform**

The microfluidic platform was fabricated using polydimethylsiloxane (PDMS) with soft lithography. The platform outline was designed using AutoCad (AutoDesk) and patterned on a silicon wafer (University Wafers) using SU-8 2050 (MicroChem). The coated wafer was then exposed to UV light through the transparency mask, which resulted in crosslinking of the photoresist imprinting the design on the wafer. A 10:1 solution of silicon elastomer base and curing agent (Ellsworth Adhesives) was poured over the wafer, degassed, and cured at 65 °C overnight. The cured PDMS was peeled from the silicon master and cut into individual devices. Inlets and outlets on the side channels of the EPC were created with 1.5 mm biopsy punches (Militex). Individual devices were irreversibly bonded against a glass coverslip using plasma treatment. The microfluidic devices were baked overnight at 65 °C and UV sterilized for 30 min prior to the experiment[1].

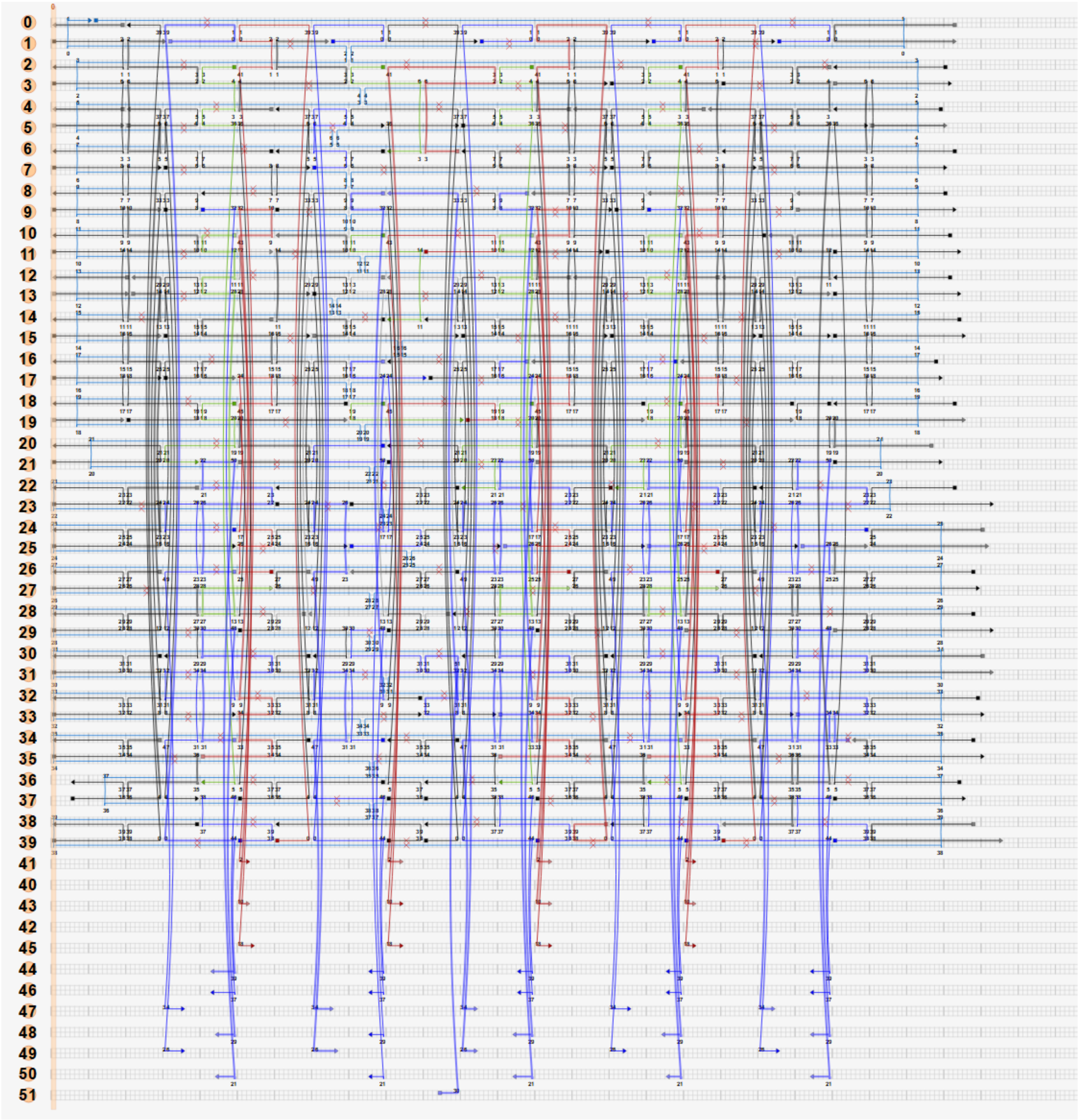

Figure S1: caDNAno design of the Cell Sensing Platform

Table S1: Custom Design Oligos for CSP

| Description and color on caDNAno | Sequence |
|---|---|
| Core (Black) | AACAATATTACCGCCTCACGCAAGTAAAGTAATTATGAAACCATCG |
| Core (Black) | AGTGAGCACATACGAGTGCTGCATCGGTGCGCGGAATACACATTCA ACCGATTGACATT |
| Core (Black) | TTAGGATTTTAGTACCTAACAGTTGATTCCCAAAACTCCTGACCCTG TAATACTTATATT |
| Core (Black) | GGCTATTAAAACAGATTTACATTGGCAGAATACCTACATTTTGACC TCAAACT |
| Core (Black) | CAACGCTAGTTGTTCCAGTTTGGTTGCGTATTGGGCGCCAGGG |
| Core (Black) | GGGAAGAACCTTATGCACAGACCAGGCGCATAGAACTGACGCAAA AGA |
| Core (Black) | ATACACTATTAAACGGCGATATATATCAGCTTCGTCCAATAAAAAG AT |
| Core (Black) | CACAAGAAAACATATAGCCAGCTTTCCGGCAC |
| Core (Black) | ATTTGGGGAAACATTAAACAGGTCAGGATTAGAAAGCGGATTGCAT CACTGCGGAA |
| Core (Black) | TAAATTGTGTCGAAATGTAACAAAGCTGCTC |
| Core (Black) | GCGGATAACGGAATAGGTGTATCTCATAGTTAGCGTAACGTAAATG ATCTTAAAC |
| Core (Black) | CAAGAAACAAACGTAAACCAGGCAAAGCGCCATCGTAAATAGCTG T |
| Core (Black) | TTCCTGTGCGCTTTCCAGTCGGGTGAGACGGGCAACAGCT |
| Core (Black) | AAACGGCTGTCTTTCCTTATCATTTCATTACCCTGCGCGT |
| Core (Black) | CAGGAGGCCGATTAACAGTGAGGCCACCGAGTTGTAGCAATACTTC |
| Core (Black) | ACGACAATATGTAGAAACCAATCAATAATGAAACGCAAAGAAACG T |
| Core (Black) | ATCTTACTCAGAGAGATAAAGCAAATCAGATATAGAAGG |
| Core (Black) | AGAAGCCTTTCAAAGCGAACCAGAAGCCCGAA |
| Core (Black) | GAGTGAGACGTCACCAGTACAAACTCAGAACCGCCACCCTGCTGA GATGGCATCA |
| Core (Black) | ACCGAAGCGATTAAGATGCGCAACTGTTGGGAATGGGATA |

| | |
|---|---|
| Core (Black) | AAATATCTTCAGTTGGGAAAACAAAATTAATT |
| Core (Black) | TGTTACTTAGCCGGAAACAAGAACCGGATATTGAGTAGTAAATTGGGCTTGAGATG |
| Core (Black) | ATATGTGAAGAGTCAATAGTGAATTTATTAGCAAGCCCAAATGGAAACAGTACAT |
| Core (Black) | TAAAACGATCATGGTCCCGTGCATCTGCCAGAATAGGAACGCCATCACAAATATTT |
| Core (Black) | TTATAATAGGGATTTGAGCACGCGCTTACCGTTTTT |
| Core (Black) | TCAACATTGACCGTAAGGGCGAAGGCGATTAGAAAAGTAAAAACAGGGAAGCG |
| Core (Black) | AAGAACGCGAGGCGTTAAGCCTTAAAAAACCG |
| Core (Black) | GGGAGAAAACCAAGTTACAATTTCTCAAACCCCAGCAAA |
| Core (Black) | TTCGCGTCGATGGGCGCATTCGCCCCCAGTCATAGCAATATTAACTGA |
| Core (Black) | CAACATTATTACAGGTCTATCATAACCCTCGTTAAATATTGGAGCCTT |
| Core (Black) | GAGATTTGCGAGGGTACAGCAGCGTTTTTCACCAGAAAACGAGACCT |
| Core (Black) | TCAGAAGCAGAGTACCAAGTTTCTGAAAAGGCTCCTCAATTTAACGG |
| Core (Black) | TTTCGGAAATTTTCAGCTGTAGCTCAACATGTTGCGGAT |
| Core (Black) | TAGCCGAAATAATAAGGCCTCTT |
| Core (Black) | TAAGAGGACCGGAAGCATTCTGCGGTTTAGCTTTTTGCTCTTGATGAT |
| Core (Black) | GAAAACTTCTACCTTAACTGATAGCCCTAAA |
| Core (Black) | TAAATCGGTTAGGTAAAGATTCAAAGAGAGATCT |
| Core (Black) | TTGAGTTATGCATGCCTGCAGGTCGACTTAGGCACTCCA |
| Core (Black) | ACAGGAGTCAAATAAGGAACCGCTTTTCATATAATCAGAGCTTTCA |
| Core (Black) | CAACAGTGCAGAAGATAGTCTTTAAAGCGTAAGACAAAACCGACCG |
| Core (Black) | TCACAATTCCACACATAACTCACATTAATTGCCTGGCCCTGA |
| Core (Black) | GCTTTGACTAGACAGGAAAGGAAGGGAAGAAATTAGAGCTTGACGGGGAACCATCA |

| | |
|---|---|
| Core (Black) | ACATCTGATGGCCAACATATTTTAATCTTCTG |
| Core (Black) | ACAAAGGAGGAAGATCCTTTAGCGTCAGACT |
| Core (Black) | ATTCTACTTCAGAGCCTCCTTTTGATAAGAGGTCTTTACCCTGACCCCCCTCAA |
| Core (Black) | TTAAATGCAATGAGAGTCTGGAGCACCCGGTTGCGGCATTT |
| Core (Black) | CGATTATAGACTAAAGTTAAAGGCCTCCAAAACATTGAATTATTATAG |
| Core (Black) | CGCCATTATCCGCGGTCACGTTGGTGTATGGCCTTCCTGTAGCCAAAAGCCCCAAAAAC |
| Core (Black) | ATTAAGAGCAGAACCGAAAGTACGGTGTCTGGTTTAATTGATAAAGC |
| Core (Black) | TACCTTTTTTAATAATAAAGACAGTAACGCCAA |
| Core (Black) | CAAGGCCGGACACCACGGAATAAGATACATAAAGGTGGC |
| Core (Black) | ACCTAAATATATGCGTCTCAACAGGACAAAAGATTAACCGTAAAAGAG |
| Core (Black) | AAGGAATTACGAGGAAACAGTTGTTGAAAA |
| Core (Black) | GAACGCGCGAAAAATAATATCCCAAACCAAGTACCGCACCAAGCAAGATGCGCCG |
| Core (Black) | GCTTAATCTAAAGCATCAATGACCATAAATC |
| Core (Black) | CCAGAACCCTAGCTGATAAATTAATGCCGGAGAAACGTTACCGTAAT |
| Core (Black) | ATCGGCCTTTTGATTAGTAATAAGAGAATAT |
| Core (Black) | AATCCTTTGCCCGAACGTATTAGACTTTACAGTTATCTA |
| Core (Black) | CACCGACTAAAGACAAAAGGGCGCCAAAAGAACTGGCATCCTTTTTAAAGTTGGGTAA |
| Core (Black) | TGAGAGATTTCAAATAGAGATAGAACCCTTTCACACGA |
| Core (Black) | TTTTGCTATTCCACAGACAGCCCACCGTACTCAGGAGGTAGCGGGGATATTTTC |
| Core (Black) | AAATCTACGTTAATAAAAAACCAAAATAGCGCTGGATAGGCTTTCGAG |
| Core (Black) | AATCATAAGGGAACCGGCTGGCTGACCTTCATCAACTTTAATCATTGTGAATTA |
| Core (Black) | CGTAGGAACCAAGAAGCTTTCCTCGTTAGAAGAGCTAAA |

| | |
|---|---|
| Core (Black) | CGCTTCTGGCCAAGCTAGCCCAATAATTGAGCGCTAATA |
| Core (Black) | AGCAAAATCAATAGAAAATTCATTTACGCAGTATGTTAGCAATGAAACGACGTTG |
| Core (Black) | GCCGGAAGGGAACAAACGGCGGATTAAATGTGAGCGAGTATGTCAATCATATGTAC |
| Core (Black) | TTGTTTGGATTATACAAAGAAACCACCAGAATTTTAAAAGTTTGAGTAACATTA |
| Core (Black) | AACCACCAAGCGGGCGCTAGGGCGCTGGCAAACTAAATCGGAACCCTGAGGTGCCGTA |
| Core (Black) | TCGTCGCTCATAGCGATAGCTTACCGGCTTAGGTTGGGTCAATCGCAAGAATAC |
| Core (Black) | TACAGTTCATATAATGGGACAAAATCATAGGTC |
| Core (Black) | CGTCAATAGATAATACCTAATAGATTAGAGCCTCAAATAATTTGAAT |
| Core (Black) | CCGAACGAACCACCAGCCACGCTGAGAGCCAGTCAATCAATATCTGGTTAGGAGC |
| Core (Black) | GGTCAGTGCAGGTCAGGAACCGCCAAGTTTGTGTATAAGAAAATAA |
| Core (Black) | AACAAGAGAATCGATGATTTTTAGAACCCTCATTTGCGGGAATAACCTAACGAGTA |
| Core (Black) | GGATTCGCCATCGGAATATCATCGCCAAGGGTTAGAACCTACCATAT |
| Core (Black) | GTAAAACAGAAATAAAATGAATAATTCATTTAACATCAACAAATCAAACCGCCTG |
| Core (Black) | TGAAAAATTGCGCCATTAAAAATACCAGTAATAAAAGGGAGCCATTGCAACAGGA |
| Core (Black) | TGTGATAACATAATTATATTTAACTCGAGCCAGTAATAACAAGTGTTT |
| Core (Black) | ACCCTCCCTATTATAACATCCAATAAATCA |
| Core (Black) | AGACTTCATAGTAAAATGTTTAGAAGAGGCTTTTGCAAAAAGGACGTT |
| Core (Black) | AAAAATCAGGTCATTTTTTTAAATTAGCATTTCTGAAACCCGTATAA |
| Core (Black) | GCCTCAGGAATTTTTGTTAAATCAGCTCATTTTTTAACCTTTGAGGG |
| Core (Black) | TGCGAATATAGGAACCCATGTACAGAGCCACC |
| Internals before TOH (Green) | CGTGCCAGTTCGTAACGGCCAGTGTGCCGGAGAAAATAC |
| Internals before TOH (Green) | TCACTGCCTGAAATTGGGGTTTATTCAGGCCTCCTTA |

| | |
|---|---|
| Internals before TOH (Green) | CCCCCGATGCGAAAGGCACCCGCTATAACGTCGGGTATT |
| Internals before TOH (Green) | CGCGGGGAGTGCTAGAGGATCCCCCAGTATCG |
| Internals before TOH (Green) | AAAGGCCGATAAAGCCAATAGTAGATGCAACTCCACCCTCCGTAACACTGAGTTT |
| Internals before TOH (Green) | GTAATGTGTGTACCAACGCGAGCATTCCATAGCCACCCTACAACGCCTGTAGCA |
| Internals before TOH (Green) | CATGGAATTCACCAGCTGACCTGAATGCGCGTTTAACCTGATTAAGACGCTGAGA |
| Internals before TOH (Green) | ATGATATTACAGCAAGGCAAAGAAGGCTTAGA |
| Internals before TOH (Green) | TCATCAGTCCAGAACCATTACCCAAATCAACCCGCGACGTACAACG |
| Internals before TOH (Green) | AACGGAAGTTTAATTTCAAGAGTAATCTTGCGAGGCGCCCCCCAG |
| Internals before TOH (Green) | ATTTAGAAGTTATTAAGGAGCGGAATTATCAATCAATAT |
| Internals before TOH (Green) | AACTAATGTCAGTGAATAAGGCTTGCCCTGACG |
| Bottom Over Hangs (Purple) | GATTGCCGCAAAATCCCTTATATTACAAAATAAACAGC CTC TGG TTA ACG TGT CTG GGC |
| Bottom Over Hangs (Purple) | GAGAGTTGCAGCAAAATCCTGTTTGATAATCCAAATAAGAAAC CTC TGG TTA ACG TGT CTG GGC |
| Bottom Over Hangs (Purple) | CCCAAATCAAGTTTTCAACGTCAAAGGGCGAATCAAGATTAGTTG CTC TGG TTA ACG TGT CTG GGC |
| Bottom Over Hangs (Purple) | AAGCGAGGCGGTAACAAGAGTCCACTATTACAATTTTATCCTGA CTC TGG TTA ACG TGT CTG GGC |
| Bottom Over Hangs (Purple) | TGGTTTTTCTTCGAGATAGGGTTGAGTACGAGCGTCTTTCCAG CTC TGG TTA ACG TGT CTG GGC |
| Bottom Over Hangs (Purple) | ATGCTTTCATAGTAAGCGGGATCGTCACCCTGCAACGGCAAAGGAAT CTC TGG TTA ACG TGT CTG GGC |
| Bottom Over Hangs (Purple) | TCGTCATTACCAGATGAGGCTTGCAGGGAGACTTTTTCTTCAGCG CTC TGG TTA ACG TGT CTG GGC |
| Bottom Over Hangs (Purple) | TGCTGAACCAGATACACTGATTGCTTTGAATCAATAACAAATCAAT CTC TGG TTA ACG TGT CTG GGC |
| Bottom Over Hangs (Purple) | ATTCATTTTGCGGAACTTCTGAATAATGGTGA CTC TGG TTA ACG TGT CTG GGC |
| Bottom Over Hangs (Purple) | AACAACCATCGCCCACGCATAACGTAAAATAGTATGGGA CTC TGG TTA ACG TGT CTG GGC |
| Bottom Over Hangs (Purple) | GCGCAGAGGCGAATTTACAGTAATCTGTAAA CTC TGG TTA ACG TGT CTG GGC |
| Bottom Over Hangs (Purple) | TAATTGTATCAACAGTATGAGGAAGTTTCCAAAACACTCATCTTTGAAGACGGTC CTC TGG TTA ACG TGT CTG GGC |
| Bottom Over Hangs (Purple) | GTGAATTATTTTCTCGTAATGCCACTACGCCTAAAACGAAAGAGCAACTTTG CTC TGG TTA ACG TGT CTG GGC |

| | |
|---|---|
| Bottom Over Hangs (Purple) | TCTCCAAGAACAACTTACAGAGGCTTTGAGCCAAGCGCGAAACAA ACTGCTCCA CTC TGG TTA ACG TGT CTG GGC |
| Bottom Over Hangs (Purple) | CCTTGCTCAGTACCTTTTACATCCAAAATTATTTGCACAATCCTGA CTC TGG TTA ACG TGT CTG GGC |
| Bottom Over Hangs (Purple) | TCGGTCATGGTGAATTTAAAGCCAGAATGGAACGTCATACATGGCT TAGTACCAG CTC TGG TTA ACG TGT CTG GGC |
| Bottom Over Hangs (Purple) | CAGTAGCGGCACCATTGACAGGAGGTTGAGGCCTTGAGTAACAGTG CATGAAAGT CTC TGG TTA ACG TGT CTG GGC |
| Bottom Over Hangs (Purple) | TTGGGAATCCTTGATATTCACAAAGTACTGGTAATAAGTGAGAAGG A CTC TGG TTA ACG TGT CTG GGC |
| Bottom Over Hangs (Purple) | AAAAGCCCCCTTCACCGGAACCAGAGCCACCACCATCCTCATATCA CCGT CTC TGG TTA ACG TGT CTG GGC |
| Bottom Over Hangs (Purple) | TGTTCAGAAGCCTGTTTAGTATCTTAATGGTTTGAAATGAACGCGA CTC TGG TTA ACG TGT CTG GGC |
| Bottom Over Hangs (Purple) | CACCACTGTTCTTACACCACCAGATGCCCCCTGCATTTCGTTACTA CTC TGG TTA ACG TGT CTG GGC |
| Bottom Over Hangs (Purple) | AGCCTAATAACAAAGTCAGAGGGTAATAAGAG CTC TGG TTA ACG TGT CTG GGC |
| Bottom Over Hangs (Purple) | CATATTATCATTAGACGGGAGAAGCTATCTT CTC TGG TTA ACG TGT CTG GGC |
| Bottom Over Hangs (Purple) | GATTTTTTACAGAGAGAATAACATAAGCAGA CTC TGG TTA ACG TGT CTG GGC |
| Bottom Over Hangs (Purple) | CTATTTTGCTTATCCGGTATTCTATTTTCAT CTC TGG TTA ACG TGT CTG GGC |
| Bottom Over Hangs (Purple) | TCTGTCCAAAGAGCGGTCAGATCCGGTCACGGCGCCCAATAGCACC CTC TGG TTA ACG TGT CTG GGC |
| Bottom Over Hangs (Purple) | AAATTGTAGGGTAGCAGCCACCACCCTCAGACCAGCATTACCATTA G CTC TGG TTA ACG TGT CTG GGC |
| Bottom Over Hangs (Purple) | TCGCATTACAACCGTTCAGTATAAAGCCAACGTATACAAATCCAGA CG CTC TGG TTA ACG TGT CTG GGC |
| Bottom Over Hangs (Purple) | CTATCAGGGAGCCGCCACCCTCAACGATTGGTAGAGCC CTC TGG TTA ACG TGT CTG GGC |
| Bottom Over Hangs (Purple) | TGCTGGTATAATTGAGAATCGCCACTAGAAACTAATGCA CTC TGG TTA ACG TGT CTG GGC |
| End Staples (Black) | TCTATCAGGGCGATTTTT |
| End Staples (Black) | TTTTT TTTGCCCCAGCAGGCGAAGCGGTCCACGCTGGTTTTT |
| End Staples (Black) | TTTTTTGGCCCACTACGTGAAAGCCGGCGTTTTT |
| End Staples (Black) | TTTTTAAGCCTGGGGTGCCTAATG |
| End Staples (Black) | TTTTTAACGTGGCGAGAACGGTACGCCTTTTT |
| End Staples (Black) | CTACAGGGCGCTTTTT |
| End Staples (Black) | TTTTTCAGCTGGCGAAAGGGGGAT |
| End Staples (Black) | TTTTTGTACTATGGTT |
| End Staples (Black) | CGCTATTACGCTTTTT |
| End Staples (Black) | TTTTTGATTCTCCGTGCATAAAGTGTATTTTT |
| End Staples (Black) | TTTTTAGAATCCTGAGATCACTTGCCTTTTTT |
| End Staples (Black) | TTTTT TAAAACTAGCAACAACCCGTCGTTTTT |
| End Staples (Black) | TTTTTGAGTAGAAGAAGCTCAATCGTCTTTTT |
| End Staples (Black) | TTTTTCAAGGATAAAAAACGGTAATCGTTTTT |

| | |
|---|---|
| End Staples (Black) | TTTTTTGAAATGGATTAGGTGAGGCGGTTTTT |
| End Staples (Black) | GTGGCACAGACATTTTT |
| End Staples (Black) | TTTTTATACATTTCGCAAATGGTC |
| End Staples (Black) | TTTTTATATTTTTGAAT |
| End Staples (Black) | GATTTAGTTTGACCATTAGTTTTT |
| End Staples (Black) | TTTTTTTAATTCGAGCTTATTTCAACGTTTTT |
| End Staples (Black) | TTTTT TCAGTATTAACCAGTTGAAAGGTTTTT |
| End Staples (Black) | TTTTTAGAGGGGGTAAAATATCGCGTTTTTTT |
| End Staples (Black) | TTTTT AATTGAGGAAGAACAATTCGAC TTTTT |
| End Staples (Black) | TTTTTTTATACCAGTCGAAGTTTTGCCTTTTT |
| End Staples (Black) | TTTTTAACTCGTATTA |
| End Staples (Black) | TTTTTATGAACGGTGTGATTTTAAGAACTGGCTCATTTTT |
| End Staples (Black) | TTTTTCAGATGATGGCAATTCTCATATTCCTGATTATTTTTT |
| End Staples (Black) | AAAGAGGACAGTTTTT |
| End Staples (Black) | TTTTTACCAAAAGGCTTTTT |
| End Staples (Black) | TTTTTTCAGGTTTAACGTCAGGAAATTGCGTAGATTTTTTTT |
| End Staples (Black) | TTTTTTTGCGCCGACAATGACAGCTTGATACCGATAGTTTTT |
| End Staples (Black) | TTTTTAGAAGATGATGAAACACAATTACCTGAGCAAATTTTT |
| End Staples (Black) | TTTTTGTCTTTCCAGACGTTAGATCTAAAGTTTTGTCTTTTT |
| End Staples (Black) | TTTTTTTAGAATCCTTGAAAAATTAATTAATTTTCCCTTTTT |
| End Staples (Black) | TTTTTTGATATAAGTATAGCCGTGCCGTCGAGAGGGTTTTTT |
| End Staples (Black) | TTTTT AATGCTGATGCAAATCTATATAACTATATGTA TTTTT |
| End Staples (Black) | TTTTT TTACCGTTCCAGTAAGAGCGCAGTCTCTGAATTTTTT |
| End Staples (Black) | TTTTTGAATAAACACCGGAATATAAGGCGTTAAATAATTTTT |
| End Staples (Black) | TTTTTCTTTTCATAATCAAAATATTAGCGTTTGCCATTTTTT |
| End Staples (Black) | TTTTTTTAGGCAGAGGCATTTAACGCCAACATGTAATTTTTT |
| End Staples (Black) | TTTTTATTGACGGAAATTATTGGGAGGGAAGGTAAATTTTTT |
| End Staples (Black) | TTTTTGATAAGTCCTGAACAACTGTTTATCAACAATATTTTT |
| End Staples (Black) | TTTTT AAACCGAGGAAACGCACAAAGTTACCAGAAGGTTTTT |
| End Staples (Black) | TTTTTGAGAATCATCTTTTT |
| End Staples (Black) | TTTTTGAAAATAGCAGCCTTTGTTTAACGTCAAAAATTTTTT |
| End Staples (Black) | TTTTTCTTGCGGGAGGTTTTGTTAGCGAACCTCCCGATTTTT |
| Top Overhangs Replacement (Red) | CACCCAGCTAAAGAACGTGGACTCTTGGGGTCAAAGGGAG |
| Top Overhangs Replacement (Red) | GACGACGAGGGTACCGAGCTCGAACTGCATTAATGAATCGGCCAA CG |
| Top Overhangs Replacement (Red) | ACACCCTGTTGCCAGAATCAAAAGAATAGCCTTCACCAGAAACCTG T |
| Top Overhangs Replacement (Red) | TTATCCCGGTGGTTCCGAAATCGCTTCACCGCGTTGCGC |
| Top Overhangs Replacement (Red) | ATCCTAATTTACGAGCAAACAACAAAAGTACCTAGGGCTATATCCA GAAAACGCT |
| Top Overhangs Replacement (Red) | TTAGCAAAATTAAGCAGAGACAGTCAAATCACCATCAAT |

| | |
|---|---|
| Top Overhangs Replacement (Red) | TTTATTTTGTCACAATCACCAGTAACAGAATCACCCTCAGTATTTTT AGGGTGAG |
| Top Overhangs Replacement (Red) | ATGGTTTACCAGCGCCTGAGCCATGTAGCGCGCTCCCTCATCATTG CCTGCCTGA |
| Top Overhangs Replacement (Red) | GTGAATAAACATTTAACAAAATCACTAACAAATTTGAGG |
| Top Overhangs Replacement (Red) | AGAAACATGAGATTTAGGAATACCACATTC |
| Top Overhangs Replacement (Red) | ATAGAAAGAAAAAAGGCGCTTTTGAGCAACAAGAAAGAT |
| Top Overhangs Replacement (Red) | AACAACTTTCGGTTTTCGGTCGCCGACGATAAACGAACT |

Table S2: Custom design oligos for detection modules

| **Cy5-labeled Detection Module on Location C3** | |
|---|---|
| Internal Cy5 Oligo _C3 | ATAGAAAGAAAAAAGGCGCTTTTGAGCAACAAGAAAGAT/Cy5 |
| Top Overhang_C3 | *CGGTCA* TAA GCG CTC ACG GAT AGG T TT TCATCAGTCCAGAACCATTACCCAAATCAACCCGCGACGTACA ACG |
| QO_C3 | Iowa Black® FQ/ ACC TAT CCG TGA GCG CTT A |
| C3_Target | ACC TAT CCG TGA GCG CTT A *TGACCG* |
| **Cy3-labeled Detection Module on Location C4** | |
| Internal Cy3 Oligo_C4 | AACAACTTTCGGTTTTCGGTCGCCGACGATAAACGAACT /Cy3 |
| Top Overhang _C4 | *CGAAGT* C ACT CCC AGG CAG CTC CAA TT AACGGAAGTTTAATTTCAAGAGTAATCTTGCGAGGCGCCCCCC AG |
| QO_C4 | Iowa Black® FQ/ TTG GAG CTG CCT GGG AGT G |
| C4_Target | TTG GAG CTG CCT GGG AGT G *ACTTCG* |

Table 3: DNA oligos used for cell functionalization

| **Cell Membrane Incorporation Oligos** | |
|---|---|
| Cholesterol Conjugated Oligo | GATGAATGGTGGGTGAGAGG /CholTEG/ |
| Bridge Oligo | CCTCTCACCCACCATTCATCTTTTTTTTT TTTTTTTTTGCCCAGACACGTTAACCAGAG |
| Fortifier Oligo | AAAAAAAAAAAAAAAAAA |

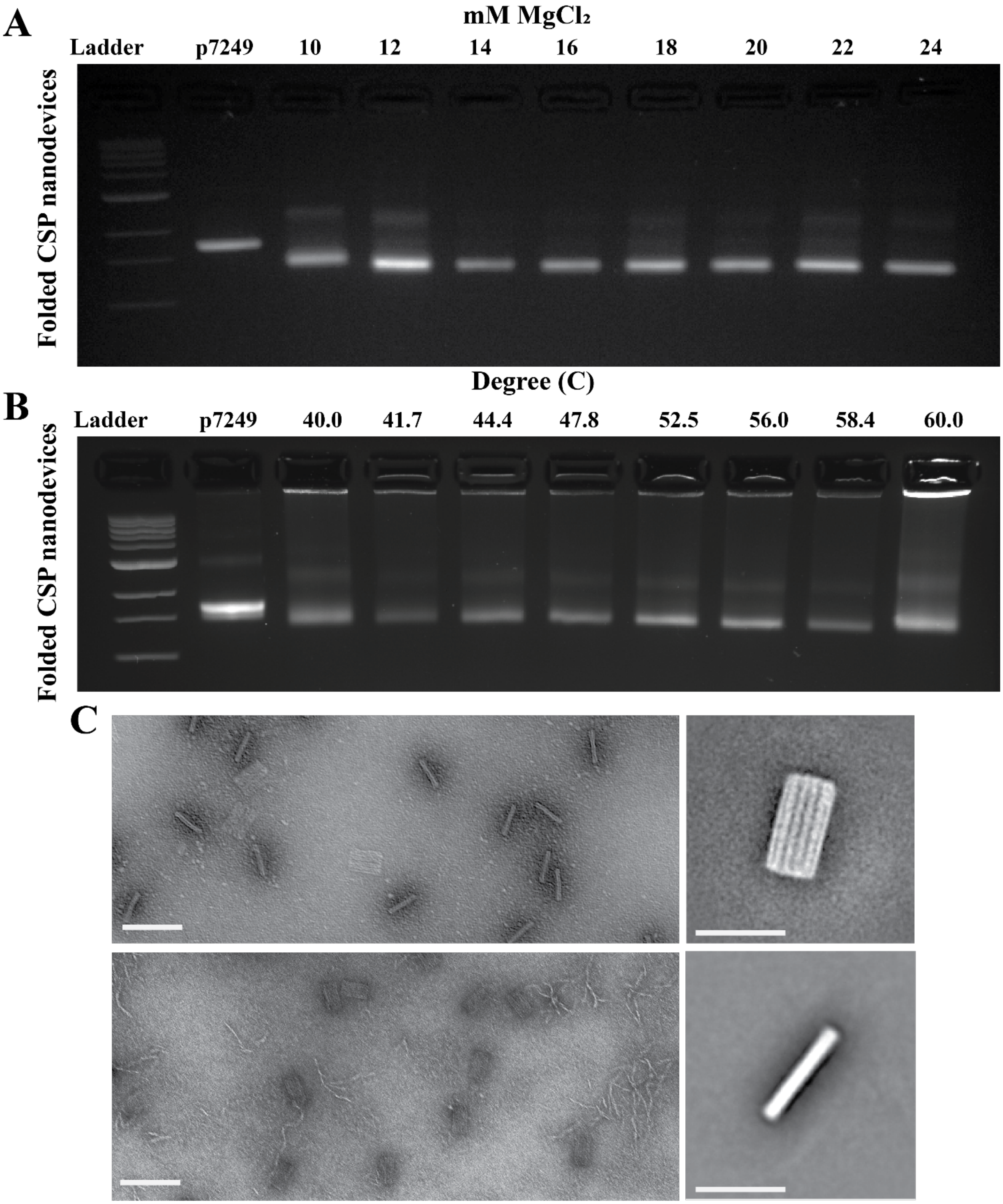


Figure S2: A. confirmation of folding of the structure using 2.5 fold protocol in different $Mg^{2+}$ concentrations (left to right, 1 kb DNA ladder (L), the 7249 M13mp18 scaffold starting material, and CSP folded in 10, 12, 14, 16, 18, 20, 22 and 24 mM $Mg^{2+}$) B. confirmation of folding of the structure using 4 hours isothermal annealing protocol (left to right, 1 kb DNA ladder, the 7249 M13mp18 scaffold starting material, and folded CSP in (40, 41.7, 44.4, 52.5, 56, 58.4, 60°C). C.

TEM confirms the successful folding of CSP in 18mM $MgCl_2$ using 2.5 day folding ramp (top left) and in 18mM $MgCl_2$ using 4 hours isothermal (52°C) ramp (Scale bar:100nm). Image averages of CSP in storage buffer using ImageJ are shown in the right. (Scale bar: 50nm)

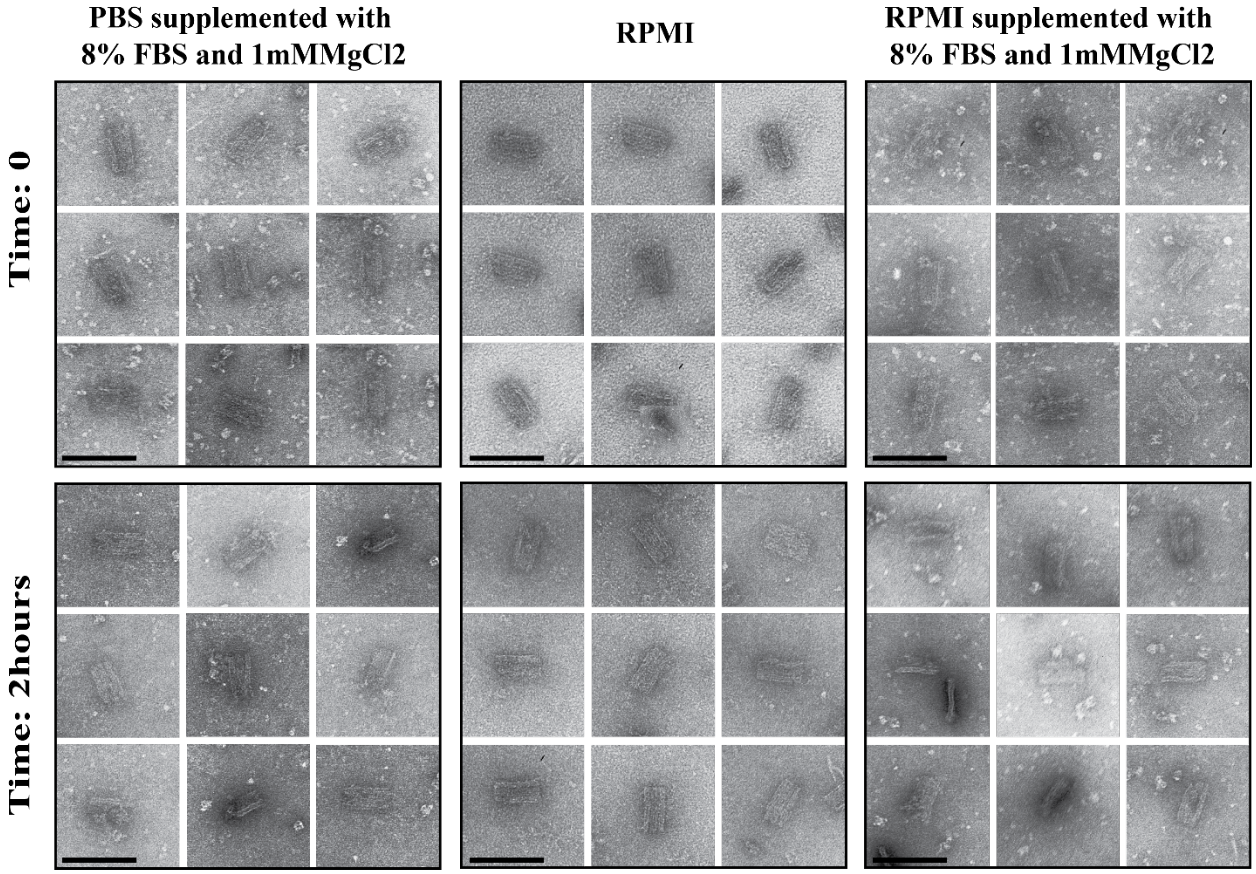


Figure S3: The stability analysis of CSP under difference cell culture media conditions using TEM. The nanostructures were incubated in each corresponding cell culture media at 37C for 4 h and then gel purified (Scale bar: 100nm).

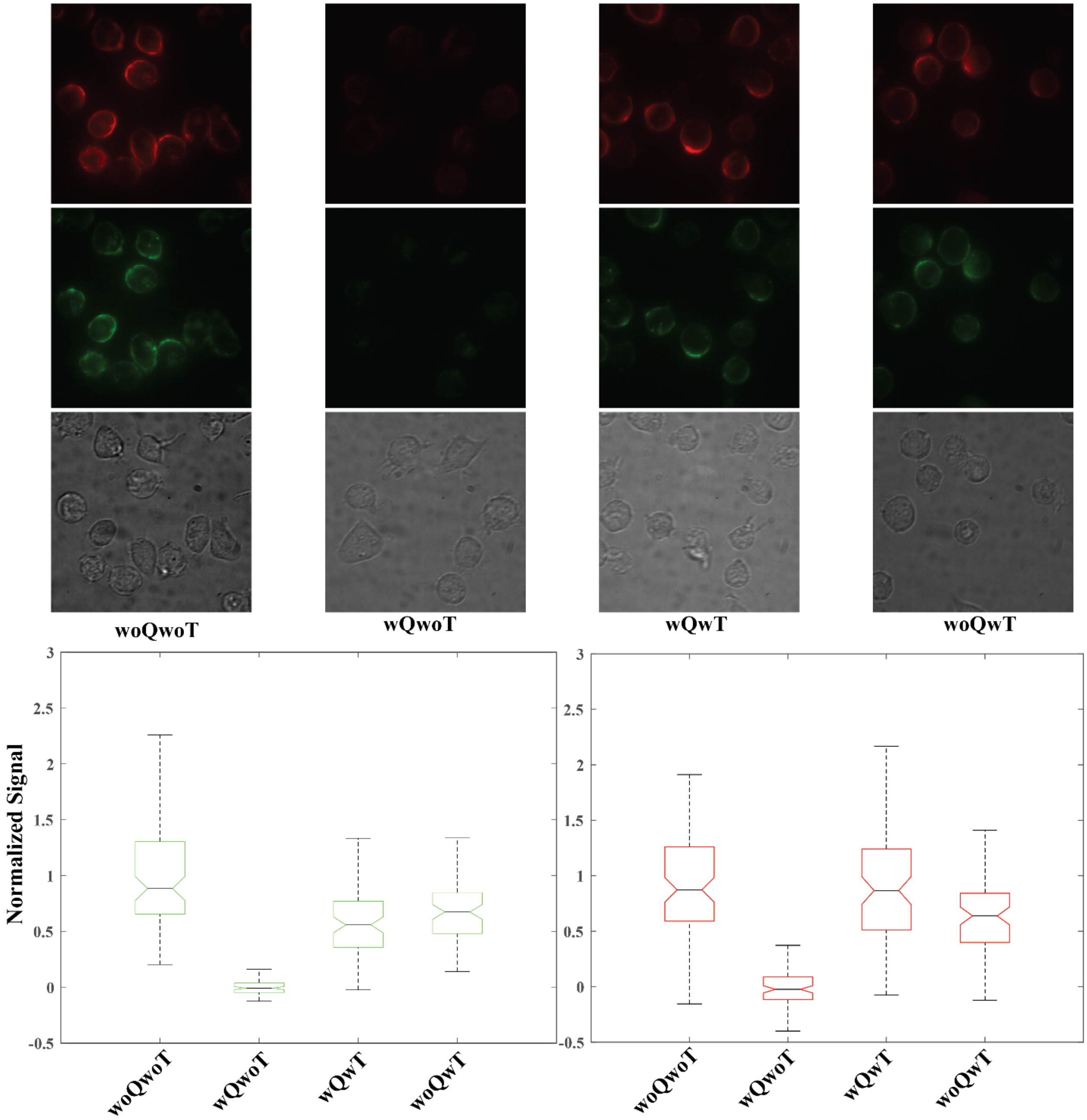


Figure S4: Detection of ssDNA targets on Dendritic cells in suspension. Top) Fluorescent and DIC images representing controls and steps taken to detect target oligos on the cell membrane. woQwoT) Dendritic cells functionalized with non-QO labeled CSP (control). wQwoT) Dendritic cells functionalized with QO labeled CSP. wQwT) wQwoT samples after the addition of both ssDNA targets. woQwT) woQwoT samples after the addition of both targets (control). Bottom) The mean fluorescence intensity attributed to the CSP bound to the surface of 80-100 single cells was quantified based on two independent experiments for each condition. The data are expressed as mean fluorescent intensity normalized relative to the average of the mean fluorescence intensity quantified for woQwoT and wQwoT conditions.

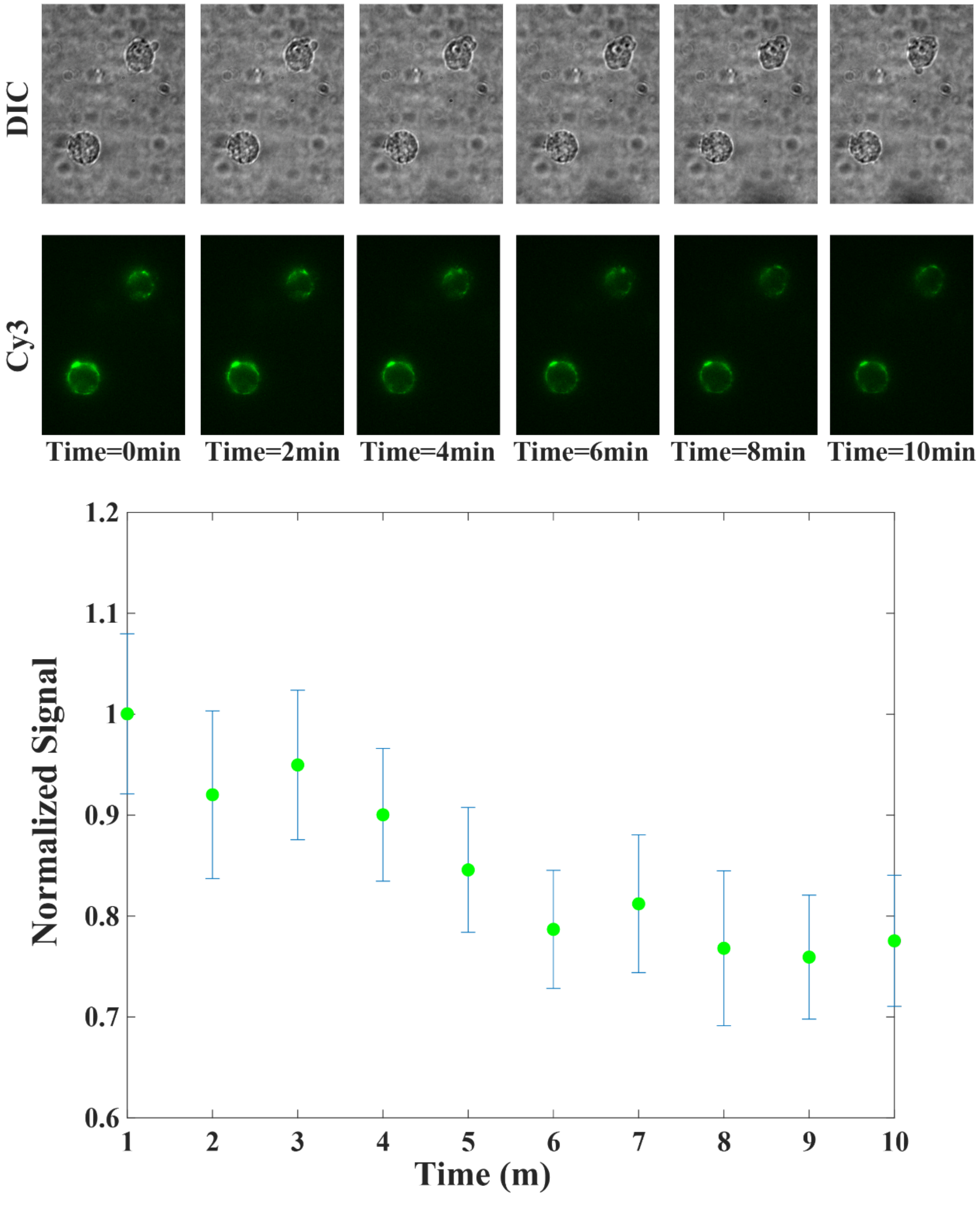


Figure S5: Photobleaching of Cy3 channel measured using cells functionalized with non-QO labeled cells. 10 Cells seeded into the collagen matrix were imaged every minute for 10 minutes with the same settings and the cell membrane signal for each cell was measured. The overall average of mean fluorescent signal was used to normalize Cy3 cell signals in Figure 3.

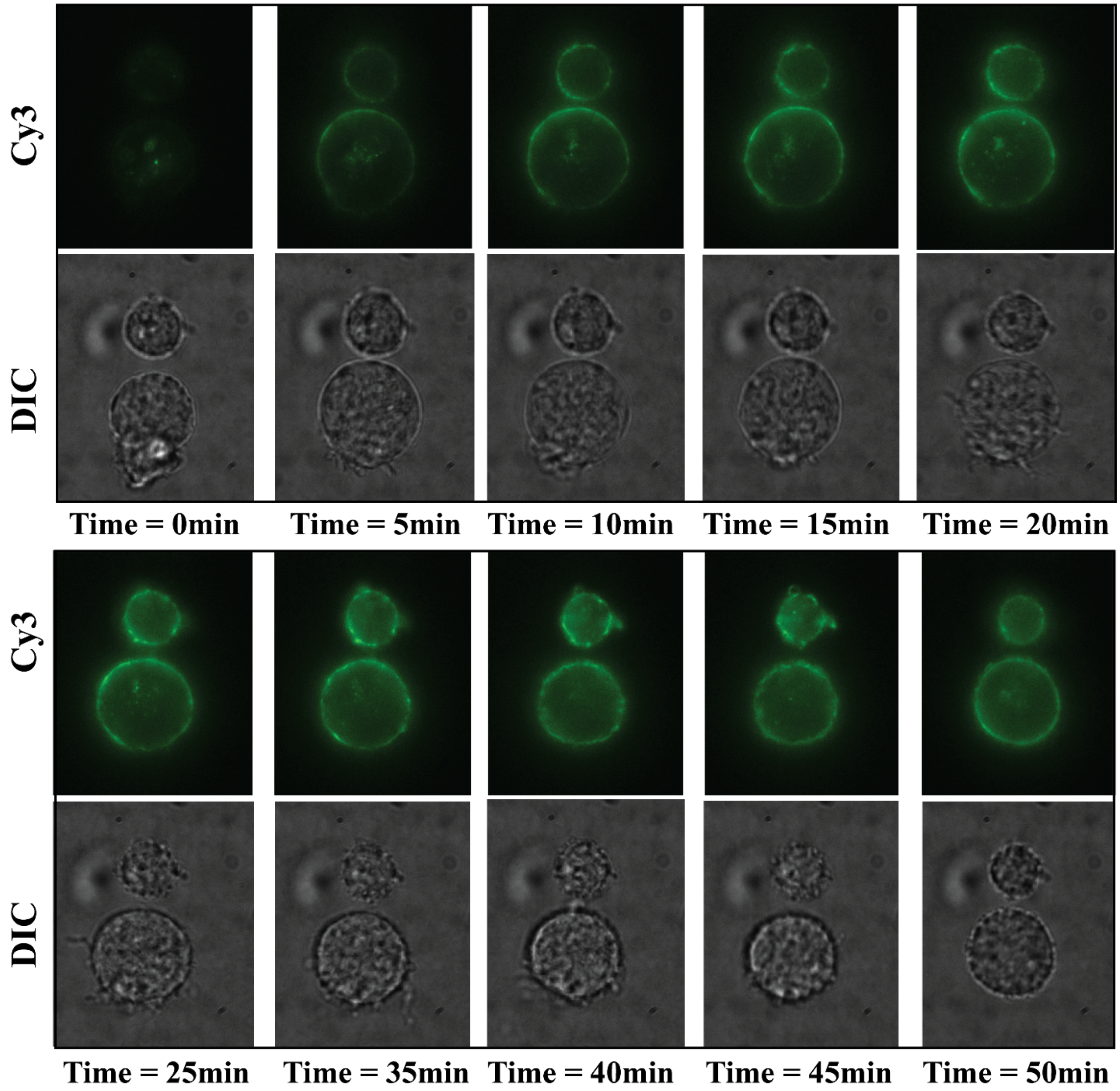


Figure S6: Long term detection of ssDNA targets on DCs in a 3-D collagen matrix using EPC. Fluorescent (Cy3) and DIC images representing the detection of target A on the cell membrane of DCs in the collagen matrix over 50min experiment.

[1] B.-H. Jo, L. M. Van Lerberghe, K. M. Motsegood, D. J. Beebe, *J. Microelectromechanical Syst.* **2000**, *9*, 76.
